# A CRISPR screening approach for identifying novel autophagy-related factors and cytoplasm-to-lysosome trafficking routes

**DOI:** 10.1101/229732

**Authors:** Christopher J Shoemaker, Tina Q Huang, Nicholas R Weir, Nicole Polyakov, Vladimir Denic

## Abstract

Selective autophagy comprises cytoplasm-to-lysosome trafficking routes that transport cargos using double-membrane vesicles (autophagosomes). Cargos are detected by receptor proteins, which typically also bind to lipid-conjugated LC3 proteins on autophagosome membranes. We dissected lysosomal delivery of four SQSTM1-like receptors by genome-wide CRISPR screening looking for novel autophagy-related (ATG) factors and trafficking routes. We uncovered new mammalian ATG factors including TMEM41B, an endoplasmic reticulum membrane protein required for autophagosome membrane expansion and/or closure. Furthermore, we found that certain receptors remain robustly targeted to the lysosome even in the absence of ATG7 or other LC3 conjugation factors. Lastly, we identified a unique genetic fingerprint behind receptor flux in *ATG7^KO^* cells, which includes factors implicated in nucleating autophagosome formation and vesicle trafficking factors. Our work uncovers new ATG factors, reveals a malleable network of autophagy receptor genetic interactions, and provides a valuable resource (<u>http://crispr.deniclab.com</u>) for further mining of novel autophagy mechanisms.

## Introduction

Macroautophagy (hereafter autophagy) is an umbrella term for a series of vesicle formation and transport mechanisms that deliver cytoplasmic material to the lysosome for degradation (Bento et al., 2016; Mizushima et al., 2011; Nakatogawa et al., 2009). During bulk autophagy, regions of cytoplasm are nonspecifically encapsulated into de novo-generated, double-membrane vesicles. These vesicles (autophagosomes) are then trafficked to lysosomes where, upon autophagosome-lysosome fusion, the inner autophagosomal membrane and its contents are degraded by lysosomal hydrolases.

There are approximately 40 autophagy-related (ATG) factors that operate at distinct stages of the autophagosome biogenesis process (Mizushima et al., 2011). Nucleation of the cup-shaped, autophagosome precursor membrane (phagophore) is thought to be mediated by the RB1CC1/ATG101/ATG13 scaffolding complex and post-Golgi vesicles containing ATG9A (Itakura and Mizushima, 2010; Kishi-Itakura et al., 2014; Koyama-Honda et al., 2013). This is followed in short succession by phagophore membrane recruitment of a phosphoinositide-3 kinase (PI3K) complex and phosphatidylinositol-3 phosphate effectors such as WIPI2 (Dooley et al., 2014). Subsequent expansion of the phagophore membrane and vesicle closure is associated with lipid conjugation of LC3-family proteins (Atg8 in yeast) via a ubiquitin-like conjugation cascade dependent on ATG7 (E1-like), ATG3 (E2-like) and ATG5/ATG12/ATG16L1 (E3-like) (Geng and Klionsky, 2008; Ichimura et al., 2000). In addition to their role in bulk autophagy, these ATG factors work together with a diverse group of proteins called autophagy receptors to enable capture of specific cytoplasmic targets, such as damaged organelles, intracellular pathogens, and protein aggregates, during autophagosome formation (Zaffagnini and Martens, 2016). In the majority of known selective autophagy cases, receptor-bound targets become tethered to growing phagophores via receptor-mediated interactions with lipidated LC3 (Stolz et al., 2014).

Defining the precise role of lipidated LC3 during either bulk or selective autophagosome formation has been historically challenging for several reasons. First, the six mammalian LC3-family proteins have similar sequences and perform related functions. A recent study overcame this issue of LC3 redundancy as it pertains to Parkin-dependent mitophagy by generating a sixtuple LC3^KO^ (LC3^NULL^) cell line (Nguyen et al., 2016). Envelopment of receptor-bound mitochondria by the phagophore membrane persisted in LC3^NULL^ cells but the resulting autophagosomes were defective for fusion with the lysosome. Second, there has been a shortage of tools for visualizing autophagosome formation without relying on microscopic reporters of LC3 conjugation. To get around this problem, another recent work monitored bulk autophagy in cells lacking LC3 conjugation factors using localization of Syntaxin 17 (STX17), a SNARE protein that is recruited to autophagosomes following vesicle closure (Itakura et al., 2012). Autophagosome-like structures (LC3 negative, Syntaxin 17 positive) formed and fused with the lysosome but failed to undergo inner autophagosomal membrane degradation, potentially because of a vesicle closure defect (Tsuboyama et al., 2016).

There are also cytoplasm-to-lysosome pathways that appear to operate normally in the absence of LC3 conjugation while still relying on other ATG factors to varying degrees. One pioneering study found conditions during which bulk protein degradation was mediated by double-membrane autophagosomes formed by an ATG5/7-independent mechanism (Nishida et al., 2009). Yet another alternative route to the lysosome was revealed in a very recent study of how the NCOA4 autophagy receptor enables iron starvation to induce degradation of ferritin (a cytoplasmic, iron-binding protein aggregate) (Goodwin et al., 2017). With the notable exception of ATG9, ferritin destruction persisted in the absence of virtually all other known ATG factors, including RB1CC1 (alternatively named FIP200), a scaffold protein essential for normal phagophore nucleation (Goodwin et al., 2017). In sum, these discoveries are beginning to re-shape the monolithic view of how ATG factors and autophagy receptors function into a more complex web of overlapping pathways for robust cytoplasm-to-lysosome delivery.

Here, we further explored the possible existence of new ATG factors and alternative routes to the lysosome by focusing on SQSTM1 and SQSTM1-like family of autophagy receptors (NDP52, TAX1BP1, and NBR1) (Birgisdottir et al., 2013). Inspired by the pH-sensitive, tandem-fluorescent reporter for monitoring LC3 lysosomal delivery (tfLC3) (Kimura et al., 2007; Pankiv et al., 2007), we used tfReceptors as phenotypic reporters for genome-wide CRISPR screening. This approach recovered virtually all known ATG factors and identified several novel candidates as screen hits. Further follow-up by quantitative microscopy and biochemical analysis revealed that TMEM41B is a previously uncharacterized endoplasmic reticulum (ER) membrane protein with a critical role in phagophore expansion and/or vesicle closure. In addition, we found that lysosomal targeting of NBR1 (and to lesser extent SQSTM1 and TAX1BP1) persisted in cells lacking LC3 conjugation factors. A subsequent screen performed using *ATG7^KO^* tfNBR1 cells enabled us to compare ATG7-dependent and ATG7-independent genetic modifiers across the entire genome. Notably, all forms of NBR1 lysosomal delivery were absolutely dependent on ATG factors implicated in phagophore nucleation (RB1CC1, ATG101, and ATG9A) but ATG7-independent delivery had a unique requirement for additional factors with known roles in alternative forms of vesicle trafficking.

## RESULTS

### Validation of tfReceptors as phenotypic reporters of autophagy

SQSTM1 and members of the SQSTM1-like family of proteins are among the most widely studied autophagy receptors in mammalian cells (Zaffagnini and Martens, 2016). We constructed six homologous gene cassettes encoding the tandem-fluorescent (tf) protein RFP-GFP by itself (tfEmpty) or as an N-terminal fusion to either LC3 (isoform B) or SQSTM1 and three SQSTM1-like family members: NBR1, TAX1BP1, and NDP52 (Fig S1A). Following cassette integration at the AAVS1 locus, we measured tf flux to the lysosome as the ratio of RFP fluorescence to GFP fluorescence, the latter of which is selectively quenched when exposed to the low pH of the lysosomal lumen (Fig 1A) (Kimura et al., 2007; Pankiv et al., 2007). As expected, inducing autophagosome formation by inhibiting mTOR with torin preferentially increased lysosomal delivery of tfSQSTM1 and tfLC3 over tfEmpty in HEK293T cells (Fig 1B). Notably, we confirmed that the torin-induced increase in the Red:Green ratio was due specifically to a decrease in GFP fluorescence as expected for this reporter system (Fig S1B). Finally, we validated that both basal and torin-induced lysosomal targeting of tfSQSTM1 was dependent on two phagophore nucleation factors, *RB1CC1* and *ATG13* (Fig 1C, Fig S1C).

**Figure 1.**
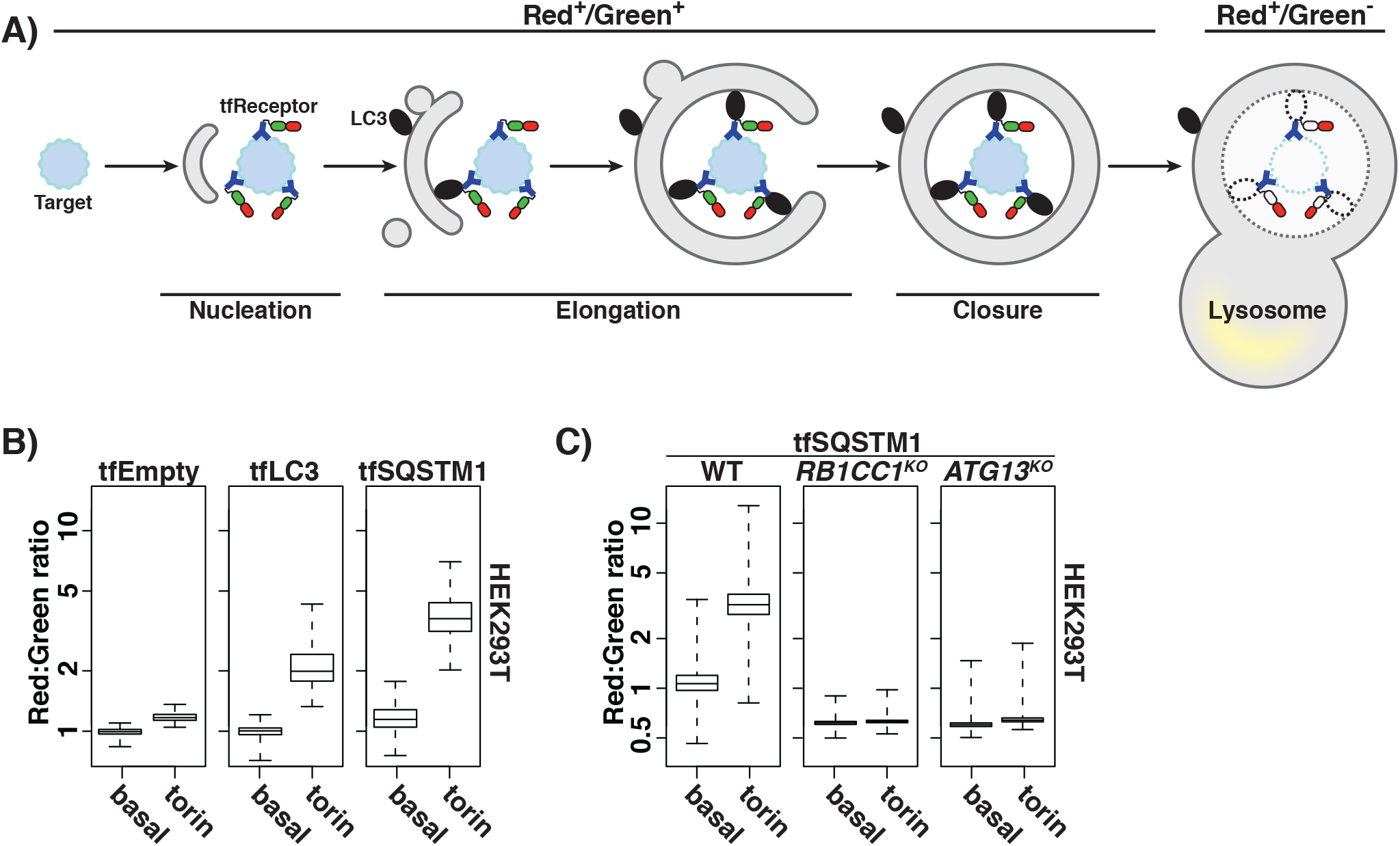
tfReceptors perform comparably to tfLC3 as reporters of autophagy. **(A)** Schematic depicting how tfReceptors measure autophagy flux. tfReceptors bind to targets and interface with remaining autophagy machinery. GFP fluorescence is selectively quenched in low pH environments, leading to an increased Red:Green ratio upon exposure of tfReceptor to the lumen of the lysosome. **(B)** HEK293T cells expressing indicated tf expression cassettes from the AAVS1 locus were analyzed by flow cytometry under basal conditions and after 18h treatment with 250nM torin. Plots show median Red:Green ratios with inner quartiles (boxed regions) and top and bottom decade indicated (whiskers). n > 10,000 cells for each sample. **(C)** Wildtype and indicated HEK293T knockout cells expressing tfSQSTM1 from the AAVS1 locus were treated and analyzed as in part B. n > 10,000 cells for each sample. See also, Figure S1.

### Genome-wide screens for tfReceptor modifiers detect known ATG factors to saturation

To get an unbiased view of autophagy pathways, we expressed our tf proteins in K562 cells (a myelogenous leukemia line), which can be cultured in suspension to facilitate genetic screening. Cells expressing either tfLC3 or tfEmpty served as receptor-agnostic controls. Bafilomycin A1 (BafA1), a known inhibitor of autophagosome-lysosome fusion, blocked basal targeting of tfLC3 and tfReceptors to the lysosome (Fig 2A). Conversely, lysosomal targeting of tfLC3 and tfReceptors was enhanced following treatment with torin (Fig 2A). Notably, tfEmpty did not respond to these treatments, implying that nonselective engulfment of tfReceptors into autophagosomes is negligible under these conditions (Fig 2A).

**Figure 2.**
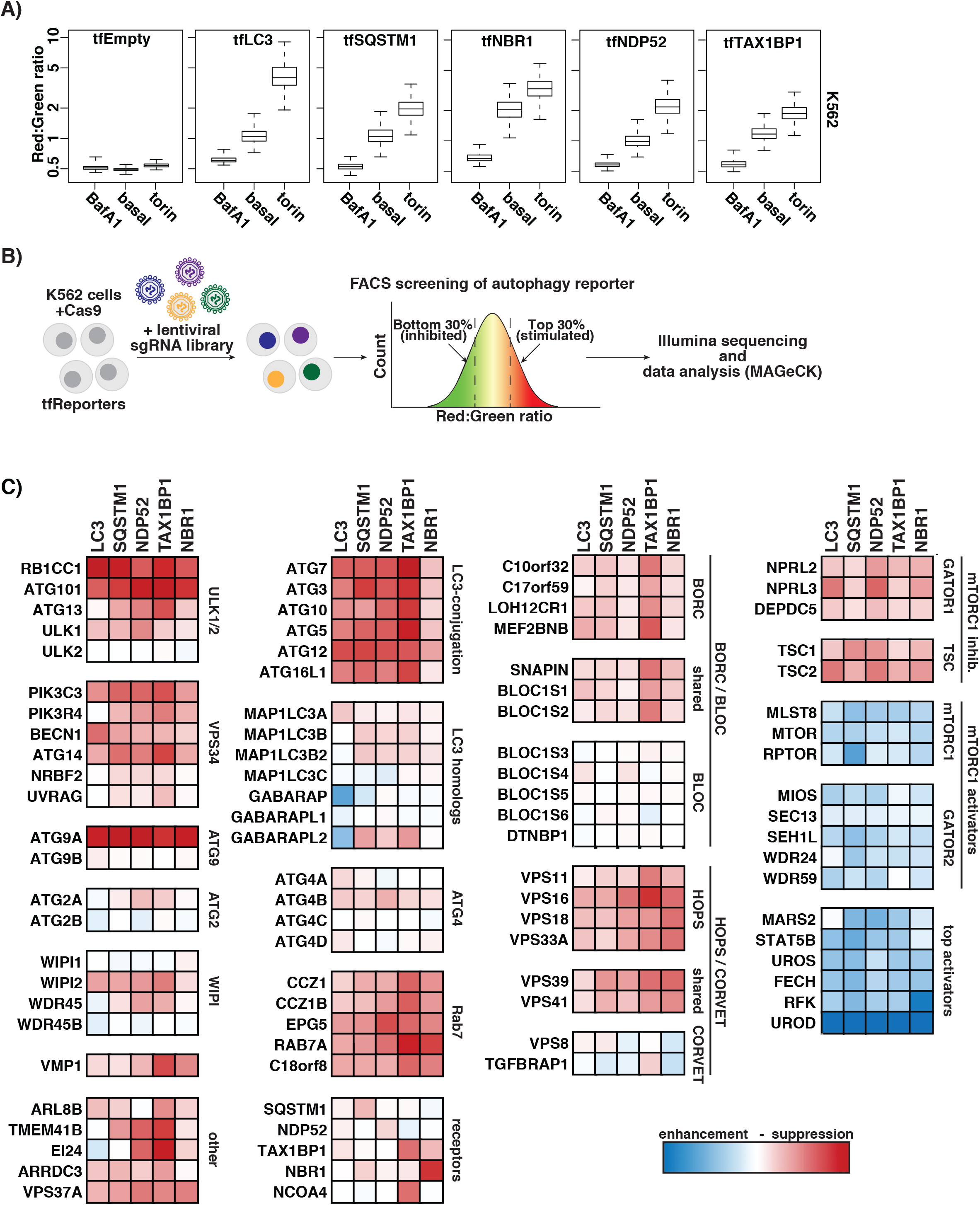
Genome-wide, pooled CRISPR screening for modifiers of tfReporters reveals known and novel autophagy-related genes. **(A)** K562 cells expressing the indicated tf proteins from the AAVS1 locus were analyzed by flow cytometry under basal conditions and after 18h treatment with 250nM torin or 100nM BafilomycinA1 (BafA1). Plots show median Red:Green ratios with inner quartiles (boxed regions) and top and bottom decade indicated (whiskers). n > 10,000 cells for each sample. **(B)** Schematic depicting a pooled CRISPR screening strategy for defining genetic modifiers of tfReporters (tfLC3 and tfReceptors). K562 cells expressing Cas9 (zeocin) were transduced with the Brunello lentiviral sgRNA library (puromycin). The top (stimulated tfReporter delivery to the lysosome) and bottom (inhibited delivery) 30% of the Red:Green ratio cell distribution cells were collected. sgRNA sequences from collected cells were obtained by an Illumina sequencing approach and analyzed by MAGeCK. **(C)** A heat map of beta scores for each gene and each tfReporter are shown. Beta scores for each gene-reporter pair were averaged across replicates. The maximum beta score for each tfReporter was used to normalize its heatmap values. For convenience, genes are clustered loosely based on their known function. Score extremes were color-coded from Red (autophagy suppression) to Blue (autophagy enhancement) with average scores uncolored (white).

Next, we performed genome-wide, pooled CRISPR knock-out screens in K562 cells coexpressing Cas9 with either tfLC3 or tfReceptors (Fig 2B). Following transduction with a library of lentivirus-encoded single guide RNAs (sgRNA; 19,114 genes), we used FACS to collect the top and bottom ~30% of cells ranked on the basis of their Red:Green ratios. Read counts of sgRNAs derived from each cell fraction were obtained by an established Illumina sequencing approach before being processed and analyzed using MAGeCK (Li et al., 2014b, 2015). The resulting output for each gene includes a beta score, a proxy for its enrichment as a phenotypic suppressor or enhancer (Fig 2C). An interactive interface for these data is also available online at <u>http://crispr.deniclab.com</u>.

Inspection of deletions that decrease tfLC3 lysosomal targeting revealed that we successfully recovered most known autophagy-related factors, thereby validating our experimental setup (Fig 2C). Comparing genetic modifiers across tfLC3 and tfReceptors revealed a strong phenotypic overlap comprising most well-established ATG factors, as well as positive and negative regulators of mTORC1. In addition, our analysis distinguished between related protein complex pairs based on their subunit ablation phenotypes. For example, in the case of the homotypic fusion and vacuole protein sorting (HOPS) and class C core vacuole/endosome tethering (CORVET) tethering complexes, we found evidence that only HOPS was required for autophagy. Similarly, we found that the BLOC-one-related complex (BORC) facilitated autophagy while the biogenesis of lysosome-related organelles complex (BLOC-1) was dispensable. Lastly, our analysis identified factors whose precise functions in autophagy have not been firmly defined (*e.g*. VPS37A, C18orf8), as well those that are completely uncharacterized (*e.g*. TMEM41B).

To validate the top screen hits, we transduced our panel of K562 cell lines with individual sgRNAs and measured phenotypes of single gene knockouts (Fig 3A,B, Fig S3A). This approach allowed us to eliminate rare screen hits (e.g. sgUROD) that were spurious because they modified the Red:Green ratio by changing RFP rather than GFP fluorescence (Fig S3B). The vast majority of tested sgRNAs confirmed that they were true screen hits whose phenotypic strengths as single gene deletions largely mirrored their relative beta score rankings (Fig 3B and 2C). Lastly, we controlled for potential tf-specific artifacts by detecting accumulation of endogenous LC3 and SQSTM1 in cells ablated for certain novel factors among the screen hits (Fig S3C). In sum, our genetic screening strategy robustly identified known ATG factors and enabled us to pursue new factors whose important roles in autophagy might have been overlooked.

**Figure 3.**
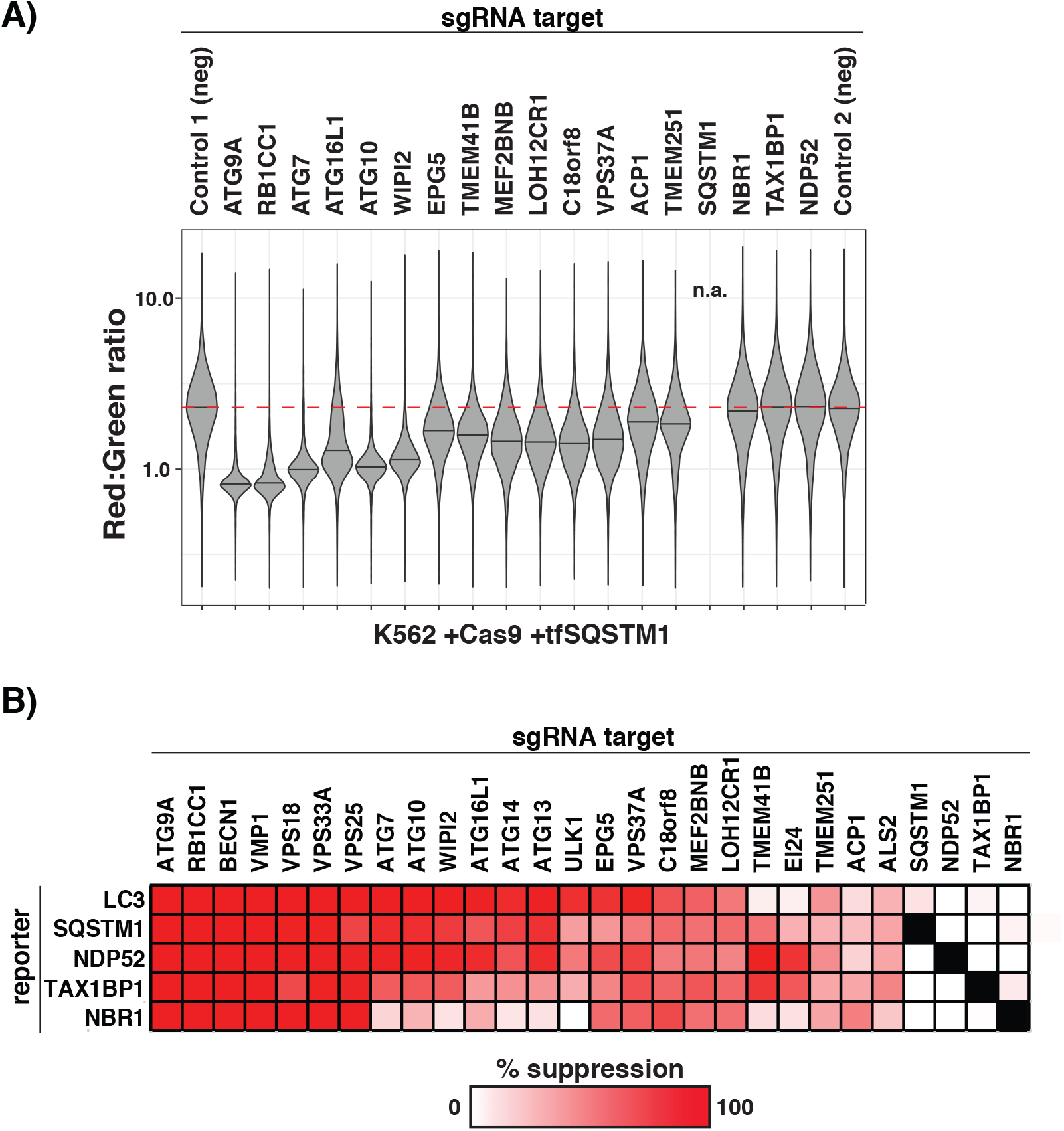
Ablation of novel screen hits robustly yields autophagy phenotypes in single gene knockout experiments. **(A)** K562 cells co-expressing Cas9 and tfSQSTM1 were transduced with individual sgRNAs against the indicated genes or with negative sgRNA controls. Cells were pre-treated with torin to maximize the dynamic range of tfSQSTM1 signals and analyzed by flow cytometry (n > 5000 cells) to determine median Red:Green ratios. Shown are violin plots generated in R using ggplot2. The median of ratios for affected cell subpopulations in each sample are indicated by a black line. Bimodal populations were deconvolved in an unbiased manner using the BifurGate function of FlowJo. Red dotted line across all samples corresponds to the median ratio for mock treated cells (Control 1). See Supplemental Figure S3A for similar data from the remaining tfReporters. **(B)** Heat map indicating the phenotypic strengths of indicated gene knockouts (as autophagy suppressors) derived from the primary data in part A and Figure S3A. The median Red:Green ratio of each population was used to calculate the fold-repression according to the following formula: (ratiosgGene-ratiosgATG9A)/(ratiosgControl-ratiosgATG9A), with the assumption that sgATG9 yields a true autophagy-null phenotype. Deeper shades of red indicate stronger suppressor phenotypes. Black boxes indicate unscored cases. See also, Figure S3.

### TMEM41B is a novel, ER membrane protein required for phagophore maturation

Despite being a complex membrane-mediated process, there is only a handful of integral membrane proteins known to be directly involved in autophagosome formation. For this reason, we focused on TMEM41B, a largely uncharacterized factor predicted to encode a multi-pass membrane protein with a C-terminal ER retention signal (Jackson et al., 1990). A global, subcellular-localization approach previously assigned TMEM41B’s residence to the ER (Itzhak et al., 2016). To confirm this finding and also to facilitate subsequent microscopy analysis of autophagosome formation in *TMEM41B^KO^* cells, we turned to adherent HEK293T cells. Using a split-fluorescent protein approach (Kamiyama et al., 2016), we tagged the N-terminus of endogenous TMEM41B (predicted to by cytosolic) with the 11^th^ beta strand of GFP (GFP11) and co-expressed a complementary GFP fragment (beta strands 1 through 10 [GFP1-10]). This resulted in the expected reticular GFP fluorescence signal that colocalized with the ER marker calnexin (Fig S4A).

Next, we made a stable *TMEM41B^KO^* cell line (Fig S4B) and established that this neither elicited an ER stress response nor diminished its induction by tunicamycin, an inhibitor of protein glycosylation in the ER (Fig S4C). Several lines of evidence argue that autophagy is severely blocked in the absence of TMEM41B. First, *TMEM41B^KO^* cells accumulated LC3 in its lipidated form (LC3-II), a phenotype that was rescued by transgenic expression of TMEM41B (Fig S4D). In addition, we found no evidence for further LC3 accumulation in *TMEM41^KO^* cells following addition of BafA1 (Fig 4A). Second, we also detected accumulation of autophagy receptors (e.g. SQSTM1, etc) in *TMEM4lB^KO^* cells (Fig 4B). Third, autophagy flux was abolished in *TMEM41B^KO^* cells as measured using tfLC3 and tfSQSTM1, which phenocopied the *ATG7^KO^* control (Fig S4E, S4F).

**Figure 4.**
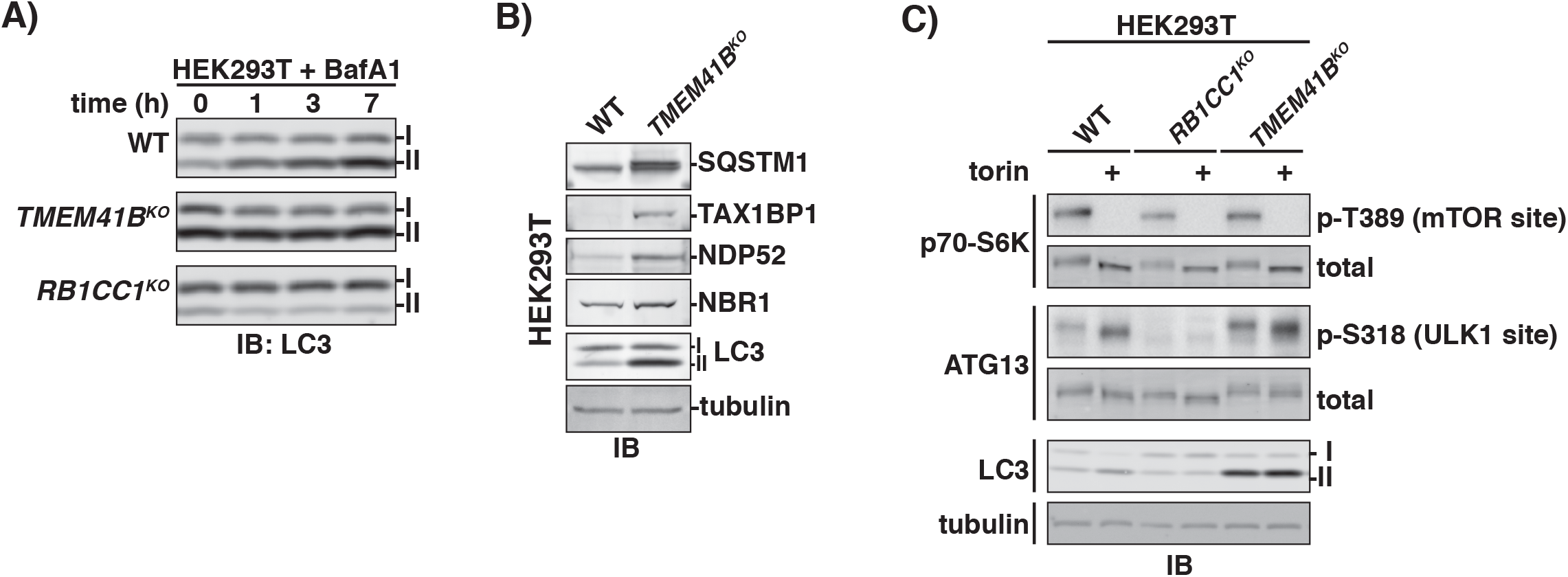
Autophagy flux is disrupted in *TMEM41B^KO^* cells despite normal induction of autophagy by torin. **(A)** Wildtype and indicated HEK293T knockout cells were treated with 250nM Bafilomycin A1 (BafA1) for the times indicated. The corresponding cell extracts were normalized by total protein and resolved by SDS-PAGE analyzed by immunoblotting (IB) for LC3. I and II indicate the unmodified and lipidated forms of LC3. **(B)** Extracts derived from wildtype and *TMEM41B^KO^* HEK293T cells were resolved by SDS-PAGE followed by immunoblotting (IB) with indicated antibodies. All samples were normalized by total protein using a BCA assay prior to loading. I and II indicate the unmodified and lipidated forms of LC3. **(C)** Wildtype and indicated HEK293T knockout cells cell were treated with 250nM torin for 3 hours or left untreated. The corresponding cell extracts were resolved by SDS-PAGE and analyzed immunoblotting (IB) with antibodies against indicated proteins or specific phosphorylation sites. All samples were normalized by total protein using a BCA assay prior to loading and even loading was verified by monitoring tubulin levels. I and II indicate the unmodified and lipidated forms of LC3. See also, Figure S4.

To define which step in autophagy requires TMEM41B, we first monitored a signal induction pathway for autophagosome initiation. mTOR kinase activity normally represses autophagy initiation by inhibiting the ULK1 kinase (Alers et al., 2012). Active ULK1 phosphorylates ATG13, a component of the RB1CC1 scaffolding complex required for phagophore nucleation (Ganley et al., 2009; Hosokawa et al., 2009; Jung et al., 2009). We found that torin treatment stimulated ATG13 phosphorylation at Ser318, a known ULK1 site, comparably in wildtype and *TMEM41B^KO^* cells (Fig 4C). For comparison, this signaling assay robustly detected loss of ULK1 activity in cells lacking RB1CC1, a known ULK1 kinase-coactivator (Fig 4C). These data argue that TMEM41B is required for a step downstream of phagophore nucleation.

To monitor phagophore elongation in *TMEM41B^KO^* cells, we visualized several markers of this process by cell microscopy. Semi-automated and custom-written scripts for image processing and segmentation allowed us to make quantitative phenotypic measurements. We found that *TMEM41B^KO^* cells displayed ~3-fold more LC3+/ SQSTM1+ punctae than wildtype cells under basal conditions (Fig 5A, 5B). To further test if these structures correspond to intermediates in autophagosome formation, we analyzed localization of the early autophagy marker WIPI2 in *TMEM41B^KO^* cells under basal and torin-induced conditions. WIPI2 transiently associates with nascent phagophores before it dissociates prior to vesicle closure (Dooley et al., 2014; Polson et al., 2010). We found that torin treatment did not further increase the level of LC3+ punctae in *TMEM41B^KO^* cells and that the majority of LC3+ structures were positive for the early autophagosome marker WIPI2 (Fig 5C, 5D). For comparison, torin induced a strong increase in LC3+ punctae formation in wildtype cells and these LC3+ structures had relatively little overlap with WIPI2 (Fig 5D). Notably, in *TMEM41B^KO^* cells both LC3+ and WIPI2+ structures were dependent on the upstream autophagy factor RB1CC1 for formation (Fig 5E, 5F, Fig S5A).

**Figure 5.**
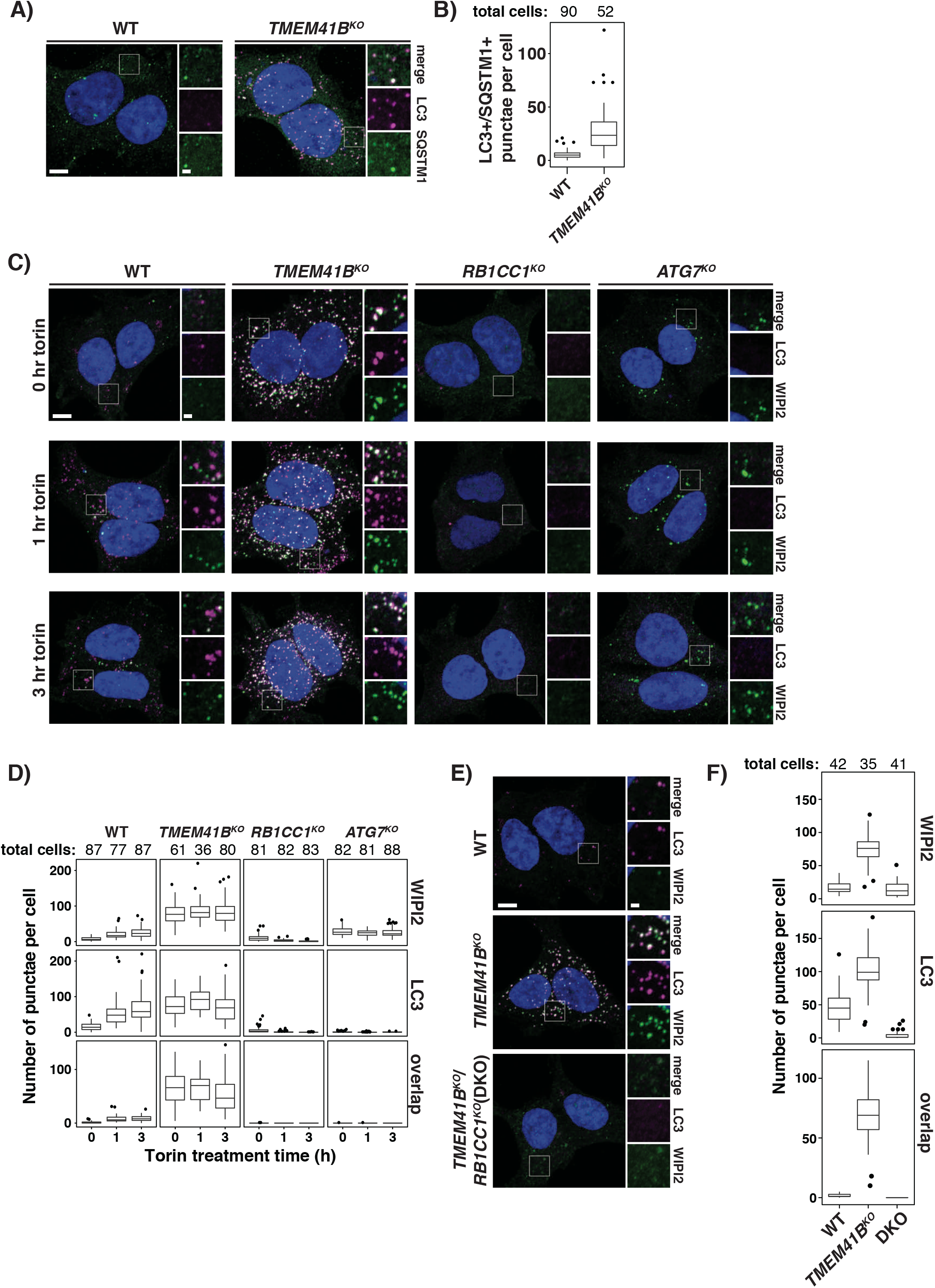
*TMEM41B^OK^* cells accumulate unresolved intermediates in autophagosome biogenesis. **(A)** Representative confocal micrographs (as maximum intensity projections) of wildtype and *TMEM41B^KO^* HEK293T cells. Selected regions (white box) of micrographs are shown as insets of single and merged channels from immunofluorescence against indicated proteins. LC3, magenta; SQSTM1, green; merged, white; Hoechst, blue. Scale bars: large panels, 5μm; small panels, 1μm. **(B)** Plots showing means of LC3+/SQSTM1+ punctae in wildtype and *TMEM41B^KO^* HEK293T cells imaged in part A (see Materials and Methods for details of quantitation) with inner quartiles (boxed regions), 1.5 interquartile ranges (whiskers), and outliers (dots) indicated. Sample size (n) for each sample is indicated. **(C)** Wildtype and indicated HEK293T knockout cells were treated with 250nM torin for the indicated times prior to analysis by confocal microscopy. Shown are representative confocal micrographs (as maximum intensity projections). Selected regions (white boxes) of micrographs are shown as insets of single and merged channels from immunofluorescence against indicated proteins. LC3, magenta; WIPI2, green; merged, white; Hoechst, blue. **(D)** Plots showing means of indicated punctae in wildtype and HEK293T knockout cells imaged in part C with inner quartiles (boxed regions), 1.5 interquartile ranges (whiskers), and outliers (dots) indicated. Sample size (n) for each sample is indicated. **(E)** Representative confocal micrographs (as maximum intensity projections) of wildtype and indicated single and double knockouts (DKO) of HEK293T cells analyzed by confocal microscopy. Selected regions (white box) of micrographs are shown as insets of single and merged channels from immunofluorescence against indicated proteins. LC3, magenta; WIPI2, green; merged, white; Hoechst, blue. Scale bars: large panels, 5μm; small panels, 1μm. **(F)** Plots showing means of indicated punctae in cells imaged in part E with inner quartiles (boxed regions), 1.5 interquartile ranges (whiskers), and outliers (dots) indicated. Sample size (n) for each sample is indicated. See also, Figure S5.

As another way of visualizing terminal steps in autophagosome formation, we turned to STX17, a SNARE protein that is recruited to mature autophagosomes following vesicle closure of elongated phagophores (Itakura et al., 2012). A recent study showed that STX17+ structures accumulate in the absence of ATG7 and we confirmed this finding (Fig S5B, S5C) (Tsuboyama et al., 2016). By contrast, we found no appreciable amounts of STX17 on LC3+ structures that accumulate in *TMEM41B^KO^* cells (Fig S5B, S5C). As a final test of whether LC3+ structures in *TMEM41B^KO^* cells represent closed autophagosomes defective for fusion with the lysosome, we carried out an LC3 protease protection assay (Fig 6A-C). To maximally accumulate any sealed autophagosomes, we inhibited autophagosome-lysosome fusion with BafA1 for 18 hours before gently lysing cells by mechanical disruption. Trypsin treatment of control wildtype cell extracts revealed that lipidated LC3 (LC3-II) was protease-resistant in the absence but not presence of Triton X-100, a non-ionic detergent that solubilizes membranes. By contrast, LC3-II was protease-sensitive in *TMEM41B^KO^* and control *RB1CC1^KO^* cells (Fig 6B, 6C). Taken together, our microscopy and biochemical evidence support the view that TMEM41B is required for a late step in phagophore maturation into fully-sealed autophagosomes.

**Figure 6.**
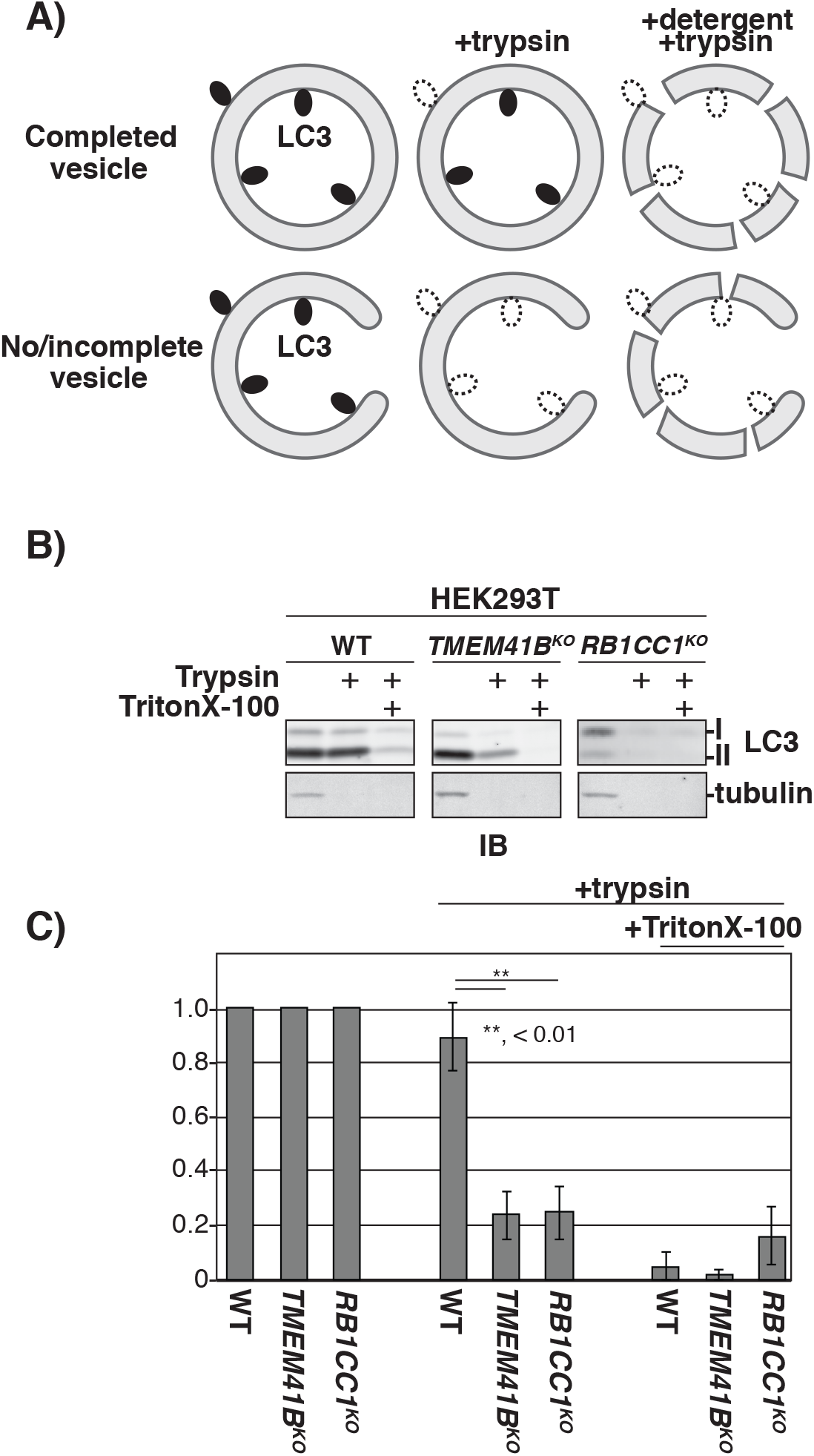
Autophagosome biogenesis in *TMEM41B^OK^* cells is arrested prior to full vesicle closure. **(A)** Schematic of protease protection assay for detecting closed autophagosomes. Lipidated LC3 (LC3-II) is indicated on the autophagosomal membrane. Dashed lines indicate proteolyzed LC3. **(B)** Wildtype and indicated HEK293T knockouts were treated for 18h with BafilomycinA1 prior to gentile, mechanical lysis. The corresponding cell extracts were treated as indicated prior to being resolved by SDS-PAGE and analyzed by immunoblotting (IB) with indicated antibodies. I and II indicate unmodified and lipidated forms of LC3. **(C)** Quantitation of protease-protection data from experiments in part B. Bar graphs show the mean +/-SD of each sample from three independent experiments. p values were determined using a student’s t-test. **,*p* < 0.01.

### Receptor-specific engagements with ATG7-independent autophagy

Returning to other results from our genetic screens, we noted that NBR1 stood out because of its unusually weak dependence on ATG7 and other LC3 lipidation factors (Fig 7A). A recent study showed that loss of RB1CC1 induces an alternative lysosome trafficking route for NCOA4 and TAX1BP1 (Goodwin et al., 2017). To examine if NBR1 can also access this route, we analyzed *RB1CC1^KO^* tfNBR1 cells but found that no evidence supporting this possibility, either under basal conditions or following torin treatment (Fig 7B, Fig S7A-C). By contrast, in *ATG7^KO^* cells lysosomal delivery of tfNBR1 was strongly stimulated by torin and inhibited by BafA1 (Fig S7C). To estimate the extent of NBR1’s engagement with ATG7-independent autophagy in K562 cells, we treated ATG9A deletions as true autophagy nulls and calculated that 65% of tfNBR1 autophagy flux was maintained in an ATG7-independent context (Fig 7C). Similarly, 30% of TAX1BP1 and 15% of SQSTM1 was delivered to the lysosome in the absence of ATG7, while delivery was negligible for NDP52 and LC3.

**Figure 7.**
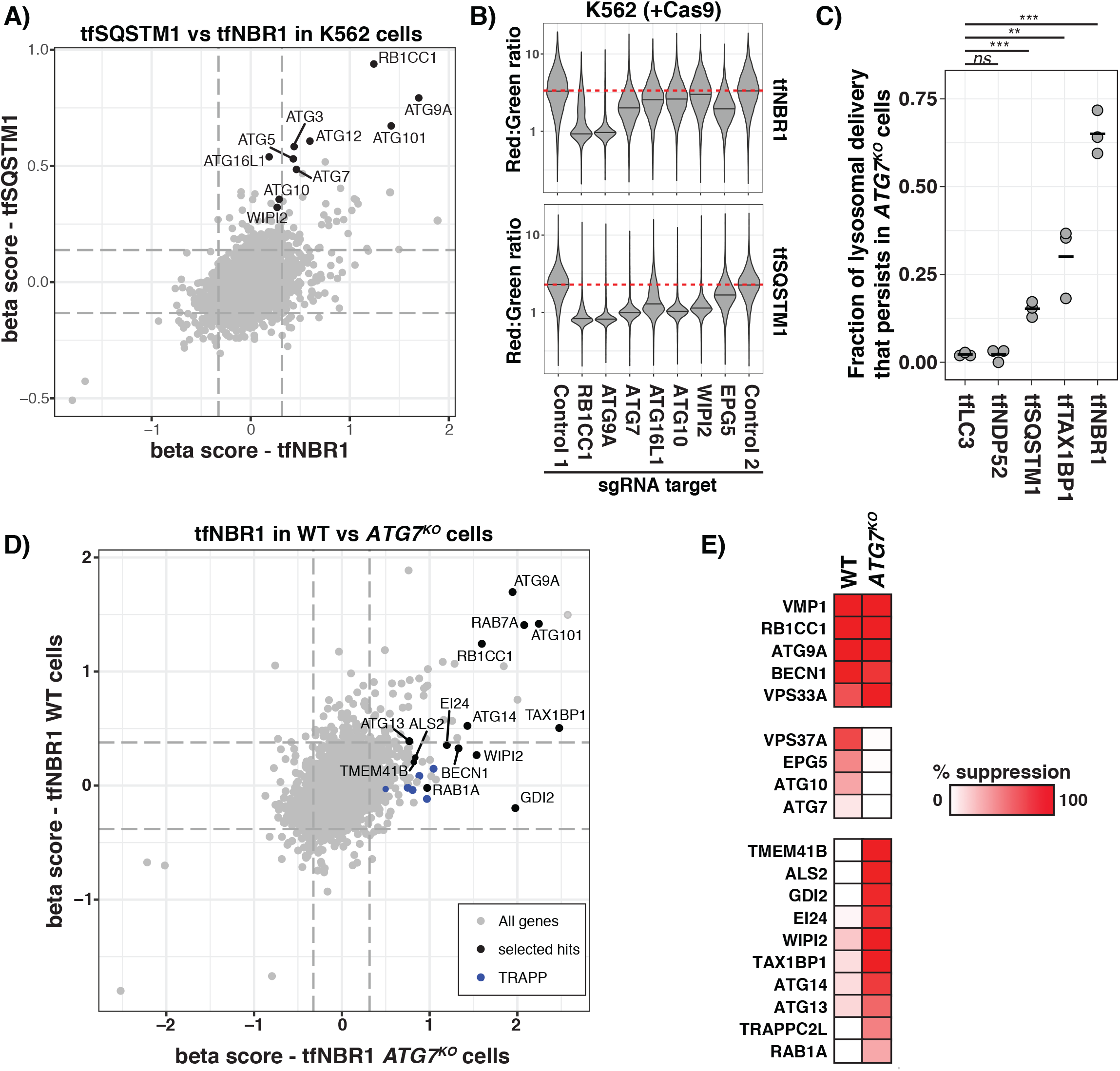
ATG7-dependent and ATG7-independent autophagy: a comparison of genetic landscapes. **(A)** Gene correlation plot (19,117 genes) of average beta scores for tfNBR1 and tfSQSTM1.: Representative autophagy-related genes are indicated in black. Dashed lines indicate top 1% of beta scores. **(B)** K562 cells co-expressing Cas9 with indicated tfReporter were transduced with individual sgRNAs against shown genes or with sgRNA controls. Cells were pre-treated with torin to I maximize the dynamic range of tfSQSTM1 signals and analyzed by flow cytometry (n>5000 cells) to determine their Red:Green ratios. Shown are violin plots generated in R using ggplot2. The median of ratios for affected cell subpopulations in each sample are indicated by a black line. Bimodal populations were deconvolved in an unbiased manner using the BifurGate function I of FlowJo. Red dotted line across all samples corresponds to the median ratio for mock treated cells (Control 1). **(C)** Dot plots of data from triplicate experiments showing the indicated tfReporter fractions delivered to the lysosome by ATG7-independent mechanisms. The median Red:Green ratio for each reporter was calculated for sgControl 1 and sgATG9A to set the upper and lower bounds, ? respectively. The median Red:Green ratio measured in sgATG7 cells was then used to calculate I the fraction of tfReporter delivered to the lysosome according to the following formula: (ratio_sgATG7_-ratio_sgATG9A_)/(ratio_sgControl1_-ratio_sgATG9A_). P-values were determined using a student’s t-test. ***,*p* <0.001; **,*p* <0.01; *ns*, not significant. **(D)** Gene correlation plot (19,117 genes) of average beta scores for tfNBR1 in wildtype and I *ATG7^KO^* cells. Notable autophagy-related genes are highlighted in black. Also highlighted (but without their gene names) are components of the TRAPP (blue) complex that scored as hits. Dashed lines, top 1% of beta scores. **(E)** Wildtype and *ATG7^KO^* K562 cells co-expressing Cas9 and tfNBR1 were transduced with individual sgRNAs for indicated genes. The median Red:Green ratio of each population was used to calculate the fold-repression according to the following formula: (ratio_sgGene_-ratio_sgATG9A_)/(ratio_sgControl_-ratio_sgATG9A_), with the assumption that sgATG9 yields a true autophagy-null phenotype. Deeper shades of red indicate stronger suppressor phenotypes. Genes were clustered on the basis of their patterns of genetic interactions.

### Forward genetic screen for modifiers of ATG7-independent autophagy

To directly probe the genetics of ATG7-independent autophagy by an unbiased approach, we performed a forward genetic screen in K562 *ATG7^KO^* cells co-expressing Cas9 and tfNBR1 (Fig 7D). From our list of screen hits and subsequent hit validation by individual sgRNA experiments, we found that lysosomal delivery of NBR1 in the absence of LC3 lipidation is absolutely dependent on several canonical ATG factors, including ATG9A, RB1CC1 and ATG101, as well as components of the late endocytic pathway including RAB7A and HOPS (Fig 7E). Next, we compared modifiers of tfNBR1 in wildtype and *ATG7^KO^* backgrounds to look for interesting genetic interactions. This analysis revealed that loss of ATG7 induced complete buffering to several ATG factors, including ATG3 and ATG10. Furthermore, we detected synthetic genetic interactions between ATG7 and several factors with poorly-defined roles in autophagy (*e.g*. EI24), as well as TMEM41B and the autophagy receptor TAX1BP1. Lastly, we found evidence that NBR1 autophagy flux in the absence of LC3 lipidation is dependent on certain vesicle trafficking (*e.g*. RAB1A and TRAPP complex components).

## Discussion

The power of forward genetics in yeast is the foundation on which the field of autophagy research firmly stands (Harding et al., 1995; Thumm et al., 1994; Tsukada and Ohsumi, 1993). Yeast genetic screens were instrumental in identifying the first autophagy receptor and the ~30 core ATG factors required for autophagosome biogenesis in yeast (Scott et al., 2001). Despite deep conservation between yeast and mammalian ATG factors, there are many mammalian-specific forms of autophagy that remain poorly understood. The recent emergence of new CRISPR/Cas9-based technologies has begun facilitating efforts to define novel mammalian autophagy factors and pathways by forward genetic screening.

Here, we combined the power of CRISPR screening with that of tandem fluorescent reporters for monitoring cytoplasm-to-lysosome targeting of several known autophagic factors. To facilitate discovery, these data are also presented online in an interactive format at <u>http://cri.spr.deniclab.com</u>. Among our screen hits, we found virtually all known ATG factors, plus autophagy modulators that only became discovered while this work was in preparation (Jia et al., 2017; Vaites et al., 2017), as well as fully uncharacterized proteins. Within this latter category, we identified TMEM41B as an ER membrane protein important for a late step in phagophore elongation and/or vesicle closure. Going forward, it will be important to test potential connections between TMEM41B and other ER-associated autophagy factors (e.g. VMP1). Another promising extension of our work will be to apply CRISPR screening to additional tf-based reporters of cytoplasm-to-lysosome targeting, including autophagy targets themselves.

Following primary screening, our focus was drawn to one receptor, NBR1, because of its unusually persistent autophagy flux in the absence of many ATG factors, including those essential for LC3 lipidation. This initial finding was surprising to us because of the well-established paradigm that NBR1 and most other autophagy receptors tether their targets to lipidated LC3 via their LC3-interacting regions (LIRs) (Birgisdottir et al., 2013). To get a better handle on this unexpected result, we carried out a secondary screen in *ATG7^KO^* tfNBR1 cells. Our specific choice for this LC3-lipidation null background was driven by two considerations. Historically, ATG7-independent autophagosomes were among the first forms of alternative autophagy to be reported (Nishida et al., 2009). Practically, *ATG7^KO^* cells are widely used in autophagy research as an autophagy-null control.

We found that genetic ablation of ATG9A, RB1CC1, and ATG101 consistently gave rise to the most severe loss-of-function phenotypes for both ATG7-dependent and ATG7-independent autophagy. In the absence of ATG7, tfNBR1 autophagy flux became increasingly dependent on certain ATG factors (e.g. ATG2B, WDR45) and less on others, including certain late endosomal factors (e.g. EPG5 and VPS37A). This type of buffering genetic interaction places LC3, EPG5, and VPS37A in the same genetic pathway. Consistently, both LC3-family proteins and EPG5 have recently been defined as vesicle/target-membrane specificity determinants in autophagosome-lysosome fusion (Nguyen et al., 2016; Wang et al., 2016). Thus, our working model is that autophagosomes formed during ATG7-independent autophagy are trafficked to lysosomes by an alternative trafficking mechanism mediated by factors that we found were selectively required for ATG7-independent but not ATG7-dependent autophagy (e.g. RAB1A, GDI2, TRAPPC2L). A mutually non-exclusive hypothesis emerges from our observation that NBR1 trafficking becomes co-dependent on TAX1BP1 in the absence of ATG7. This autophagy receptor is known to interact with myosin IV and was previously postulated to control vesicle trafficking by the actin/myosin network (Tumbarello et al., 2015). Intriguingly, a recent study showed how activation of TBK1 kinase enables TAX1BP1 to direct lysosomal targeting of ferritin by a novel ESCRT-mediated process (Goodwin et al., 2017). Our work implicates TAX1BP1 in another noncanonical cytoplasm-to-lysosome pathway and should help raise the interest level in this understudied autophagy receptor.

Mechanistically, it remains unclear how NBR1, as well as TAX1BP1 and SQSTM1 to lesser extents, engages the autophagosome biogenesis process in the absence of LC3 lipidation. In yeast, several receptors are known to locally induce autophagosome formation by binding to a C-terminal region of Atg11 (Kamber et al., 2015), which has weak homology to a C-terminal sequence of RB1CC1 (Li et al., 2014a; Yorimitsu and Klionsky, 2005). Analogous mechanisms could explain our observation that NBR1’s lysosomal delivery remains selective in the absence of ATG7 albeit in a manner still dependent on RB1CC1 and ATG101. Consistently, there is evidence that mammalian autophagy receptors (both mitophagy receptors (Lazarou et al., 2015) and TRIMs (Kimura et al., 2016)) can recruit an RB1CC1-containing complex (RB1CC1/ATG101/ATG13/ULK1) to locally induce autophagosome formation.

LC3 positive and negative double-membrane vesicles have previously been shown to coexist in certain cell types (Nishida et al., 2009). To what extent do receptors then engage alternative autophagy pathways in cells permissive for LC3 lipidation? Answering this important question will necessitate development of receptor-based reporters that differentiate flux through related pathways comprising many overlapping components. At one extreme, it is possible that LIR-containing receptors normally engage only LC3-marked autophagosomes. This would then imply that ablation of ATG7 induces compensatory mechanisms, which bypass the need for receptor-LC3 interactions. It also remains unclear what distinguishes receptors that engage ATG7-independent autophagy from those that don’t (e.g. NBR1 vs NDP52). Since NBR1 is known to interact with a diverse set of autophagy targets (Deosaran et al., 2013; Kenific et al., 2016; Kirkin et al., 2009; Lamark et al., 2009), one intriguing possibility is that its engagement with this alternative autophagy pathway is target dependent.

Our systematic, genome-wide analysis of NBR1 autophagy flux casts a new light on the roles of ATG factors within the autophagy cannon. We found that some are absolutely essential (e.g. RB1CC1 and ATG9A) whereas others (e.g. WIPI2 and ATG14) only become critical in the absence of ATG7. At a molecular level, RB1CC1 and ATG9A are among the first ATG factors to be recruited to ER-associated sites at which LC3-marked autophagosomes eventually form (Koyama-Honda et al., 2013). RB1CC1 doesn’t have a strong yeast homolog by sequence analysis, but is very likely structurally and functionally equivalent to Atg17 and Atg11 in yeast (Li et al., 2014a). Recent studies have proposed that Atg17 is sufficient to remodel Atg9 vesicles into a cupshaped structure (Bahrami et al., 2017; Ragusa et al., 2012). Our work makes it further imperative to establish if RB1CC1 scaffolding of ATG9A vesicles is a conserved phagophore nucleation mechanism. Even more speculatively, our data suggests that autophagosomes that are produced by a common phagophore nucleation process can then engage a variable set of mechanisms involved in downstream autophagy steps. Further mapping of pairwise genetic interactions between ATG factors is a promising approach for teasing apart these complexities inherent in the mammalian ATG network.

---

**Figure S1 -.**
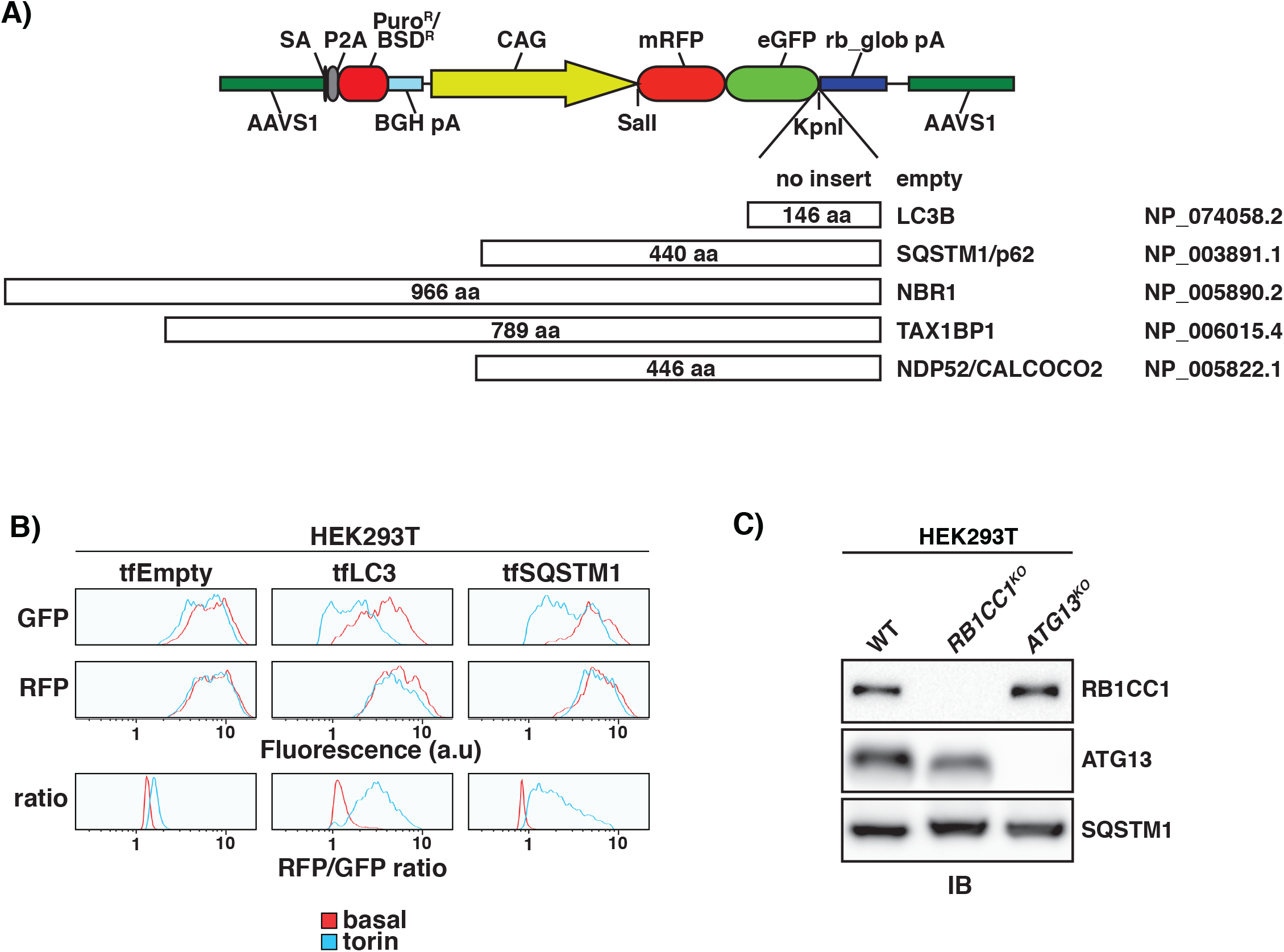
Figure 1, supplement 1. Generation and validation of tfReceptor autophagy reporters. **(A)** Diagram of gene cassette used for expressing all tf proteins in this study. Salient cassette features are color coded. Shown below are ATG factors that were expressed as N-terminal tf fusions with their length, common names, and RefSeq accession number listed left to right. SA, splice acceptor; P2A, self-cleaving peptide; Puro^R^/BSD^R^, puromycin or blasticidin resistance cassette; BGH pA, bovine growth hormone polyadenylation signal; CAG, CAG promoter sequence; AAVS1, AAVS homology arms. All inserts were positioned at the KpnI restriction site. **(B)** HEK293T cells expressing the indicated tf proteins from the AAVS locus were grown under basal conditions or treated with torin. Shown are flow cytometry traces of GFP and RFP fluorescence (arbitrary units), both as individual signals and as a ratio (Red:Green). **(C)** Extracts derived from cells with indicated genotypes were normalized by total protein levels using a BCA assay and resolved by SDS-PAGE followed by immunoblotting (IB) with indicated antibodies.

**Figure S3 -.**
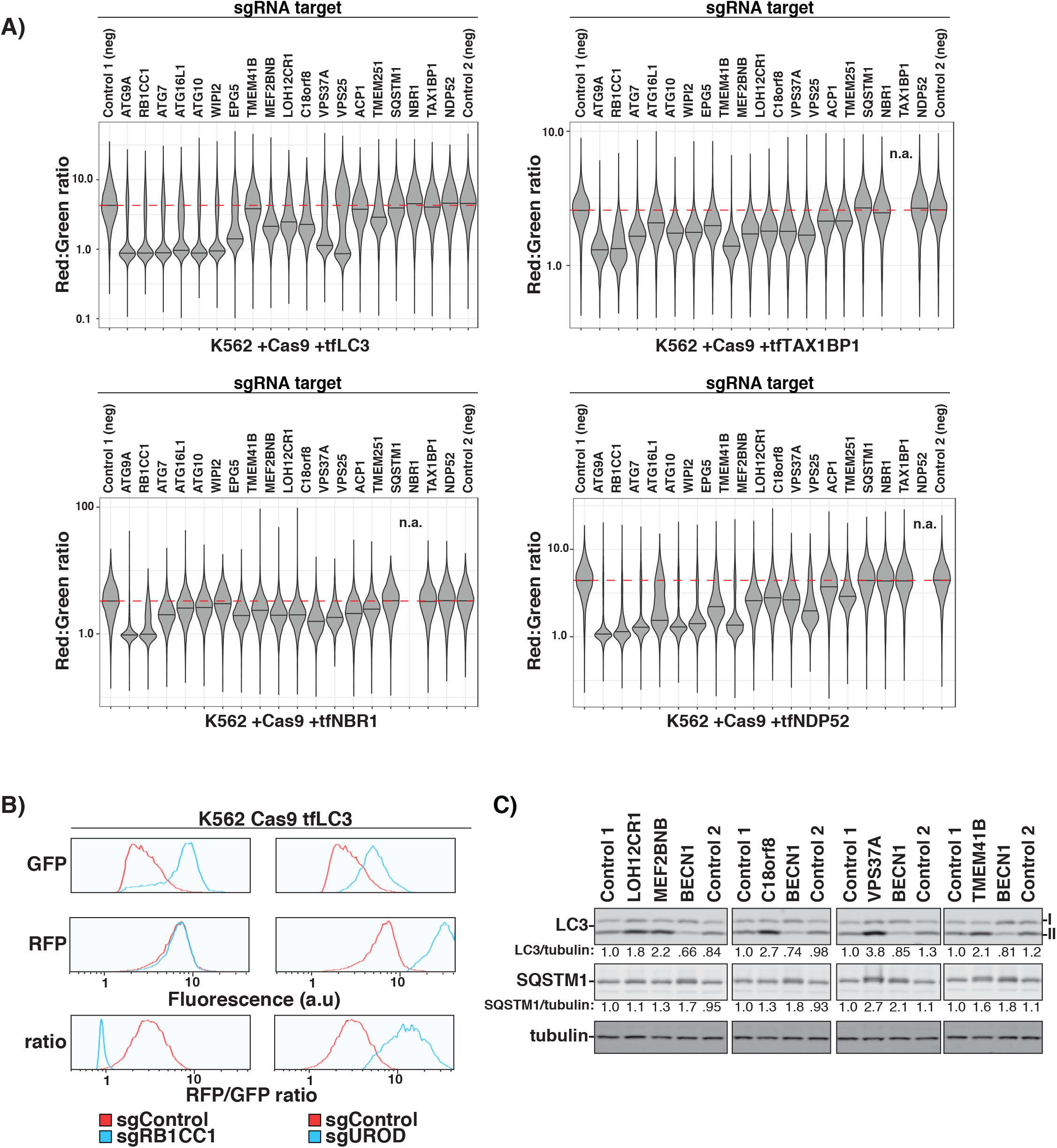
Figure 3, supplement 1. Confirmation of sgRNA effects on autophagy reporters. **(A)** K562 cells co-expressing Cas9 and indicated tfReporters were transduced with individual sgRNAs against the shown genes or with negative sgRNA controls. Cells were treated and analyzed as in Figure 3A. These data are represented as part of the heatmap in Figure 3B. n > 5000 cell each. **(B)** K562 cells co-expressing Cas9 and tfLC3 were transduced with sgRNAs against the indicated genes or with a negative sgRNA control. Shown are flow cytometry traces of GFP and RFP fluorescence (in arbitrary units), both as individual signals and as a ratio (Red:Green). Cells were treated and analyzed as in Figure 3A. **(C)** K562 cells expressing Cas9 were transduced with individual sgRNA against the indicated genes or negative sgRNA controls. The corresponding cell extracts were resolved by SDS-PAGE followed by immunoblotting (IB) with indicated antibodies. I and II indicate the unmodified and lipidated forms of LC3, respectively. Numbers indicate the indicated protein ratios and were set to 1 for the Control 1 in each boxed data set.

**Figure S4 -.**
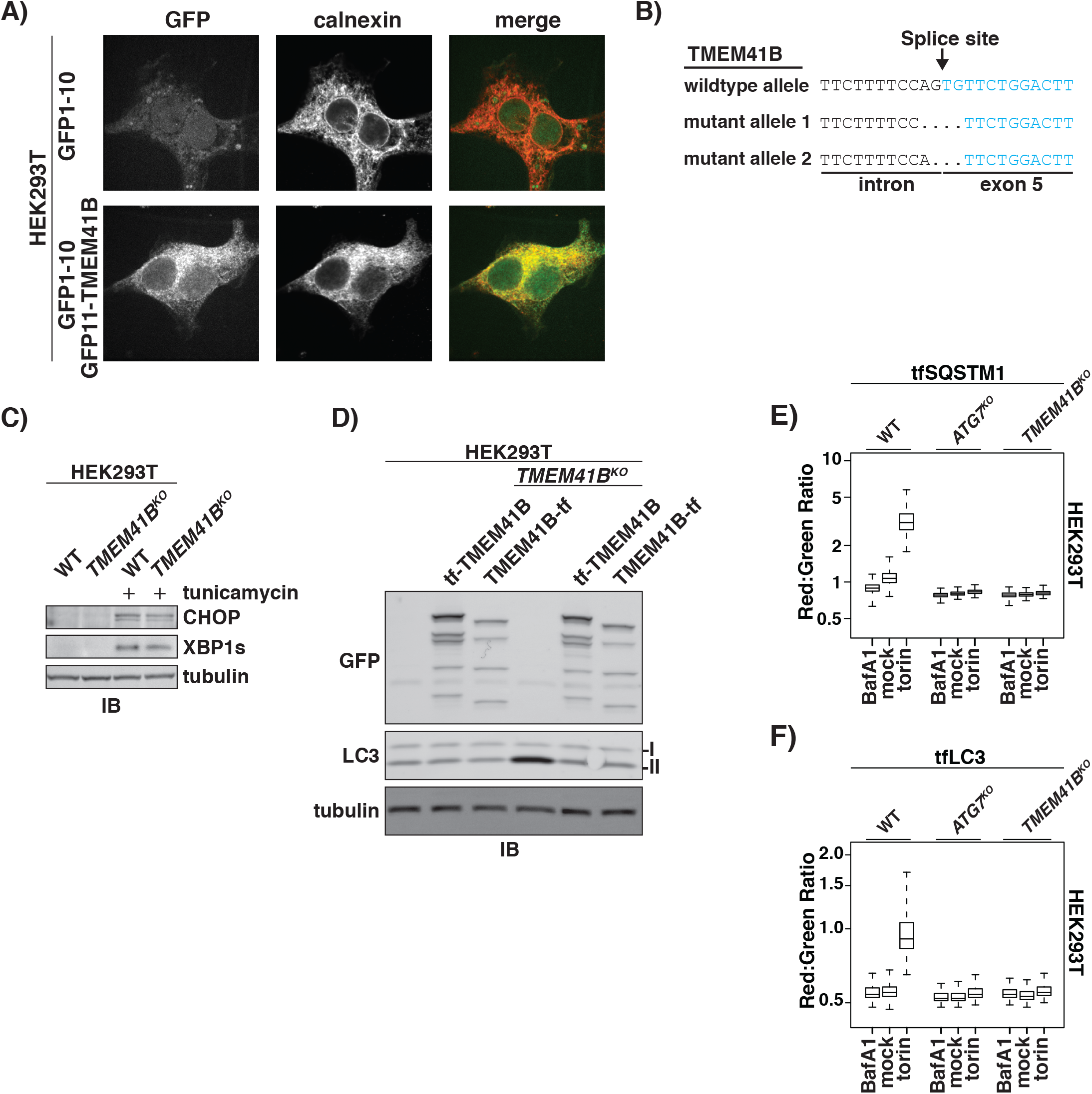
Figure 4, supplement 1. TMEM41B deletion inhibits autophagy. **(A)** Wildtype HEK293T cells (top) or expressing endogenous TMEM41B with an N-terminal GFP11 tag (bottom) were transduced with a lentivirus expressing GFP1-10 and analyzed by confocal microscopy. Shown are confocal slice micrographs of GFP fluorescence and calnexin immunofluorescence, both as individual signals and merged. **(B)** Sequence of mutant alleles in *TMEM41B^KO^* HEK293T cells. Wildtype allele with indicated splice site is shown for comparison. Color coding was used to distinguish intron and exon sequences. **(C)** Extracts derived from wildtype and *TMEM41B^KO^* HEK293T cells were normalized for total protein by BCA and resolved by SDS-PAGE followed by immunoblotting (IB) with indicated antibodies. **(D)** Extracts derived from wildtype and *TMEM41B^KO^* HEK293T cells expressing indicated tf fusions of TMEM41B were normalized for total protein by BCA resolved by SDS-PAGE followed by immunoblotting (IB) with indicated antibodies. I and II indicated unmodified and lipidated forms of LC3. **(E and F)** Wildtype and indicated HEK293T knockout cells expressing indicated tfReporters were analyzed by flow cytometry under basal conditions and after 18h treatment with 250nM torin or 100nM BafilomycinA1 (BafA1). Plots show mean Red:Green ratios with inner quartiles (boxed regions) and top and bottom decade indicated (whiskers). n > 5000 cells each sample.

**Figure S5 -.**
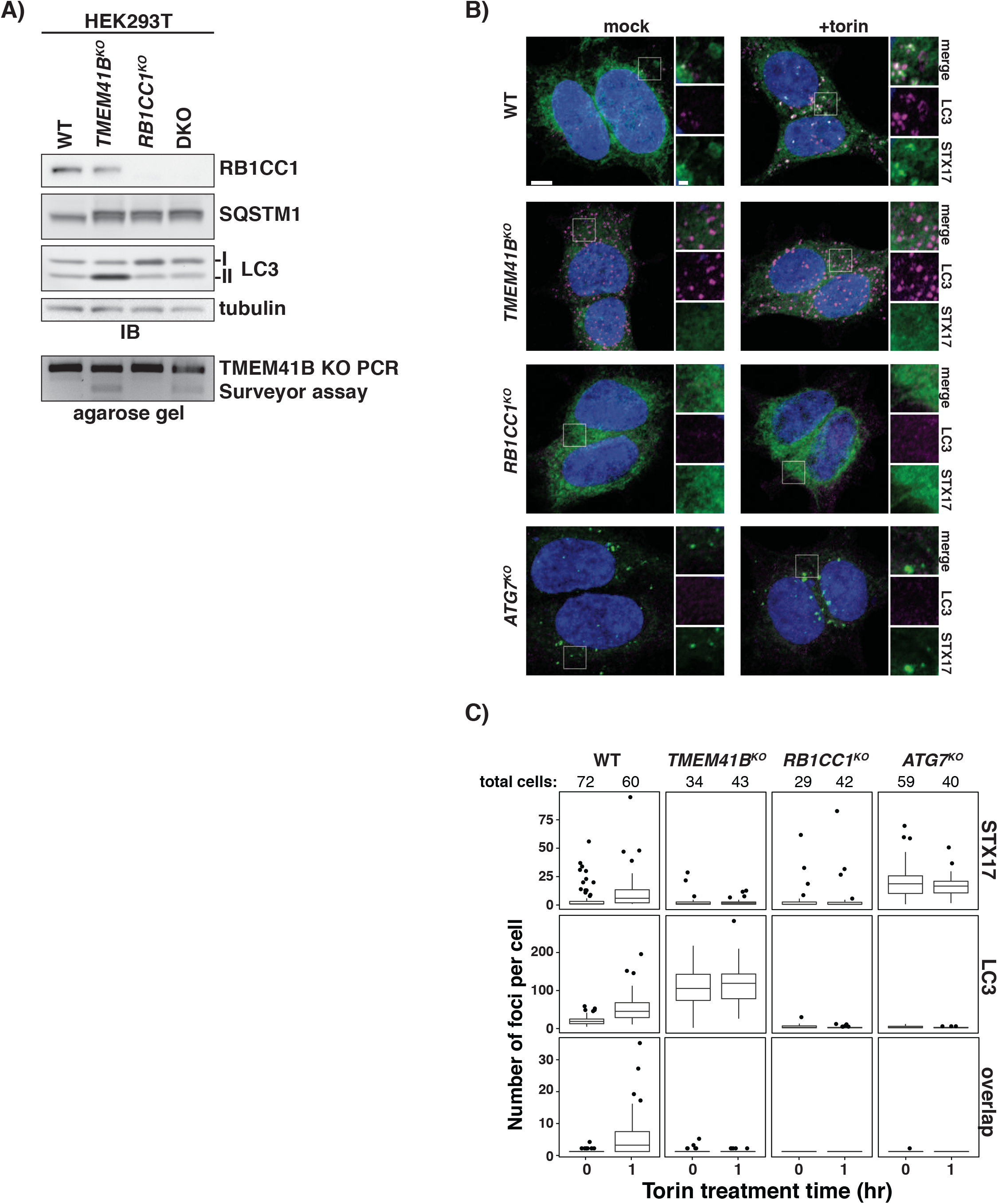
Figure 5, supplement 1. TMEM41B deletion arrests autophagy on-pathway prior to autophagosome closure. **(A)** Extracts derived from wildtype and indicated HEK293T single and double knockout (DKO; *TMEM41B^KO^IRB1CC1*^KO^) cells were resolved by SDS-PAGE and analyzed by immunoblotting (IB) for indicated proteins. I and II indicate unmodified and lipidated forms of LC3. Also shown are PCR results from a T7 endonuclease assay used to confirm TMEM41B gene knockouts. **(B)** Wildtype and indicated HEK293T knockout cells expressing GFP-STX17TM were treated with 250nM torin for one hour or left untreated (mock) prior to confocal microscopy. Shown are representative confocal micrographs (as maximum intensity projections). Selected regions (white boxes) of micrographs are shown as insets of single and merged channels from intrinsic GFP fluorescence or immunofluorescence against indicated proteins. LC3, magenta; STX17, green; merged, white; Hoechst, blue. Scale bars: large panels, 5μm; small panels, 1μm. **(C)** Plots showing means of indicated punctae in wildtype and HEK293T knockout cells imaged in part B with inner quartiles (boxed regions), 1.5 interquartile ranges (whiskers), and outliers (dots) indicated. Sample size (n) for each sample is indicated.

**Figure S7 -.**
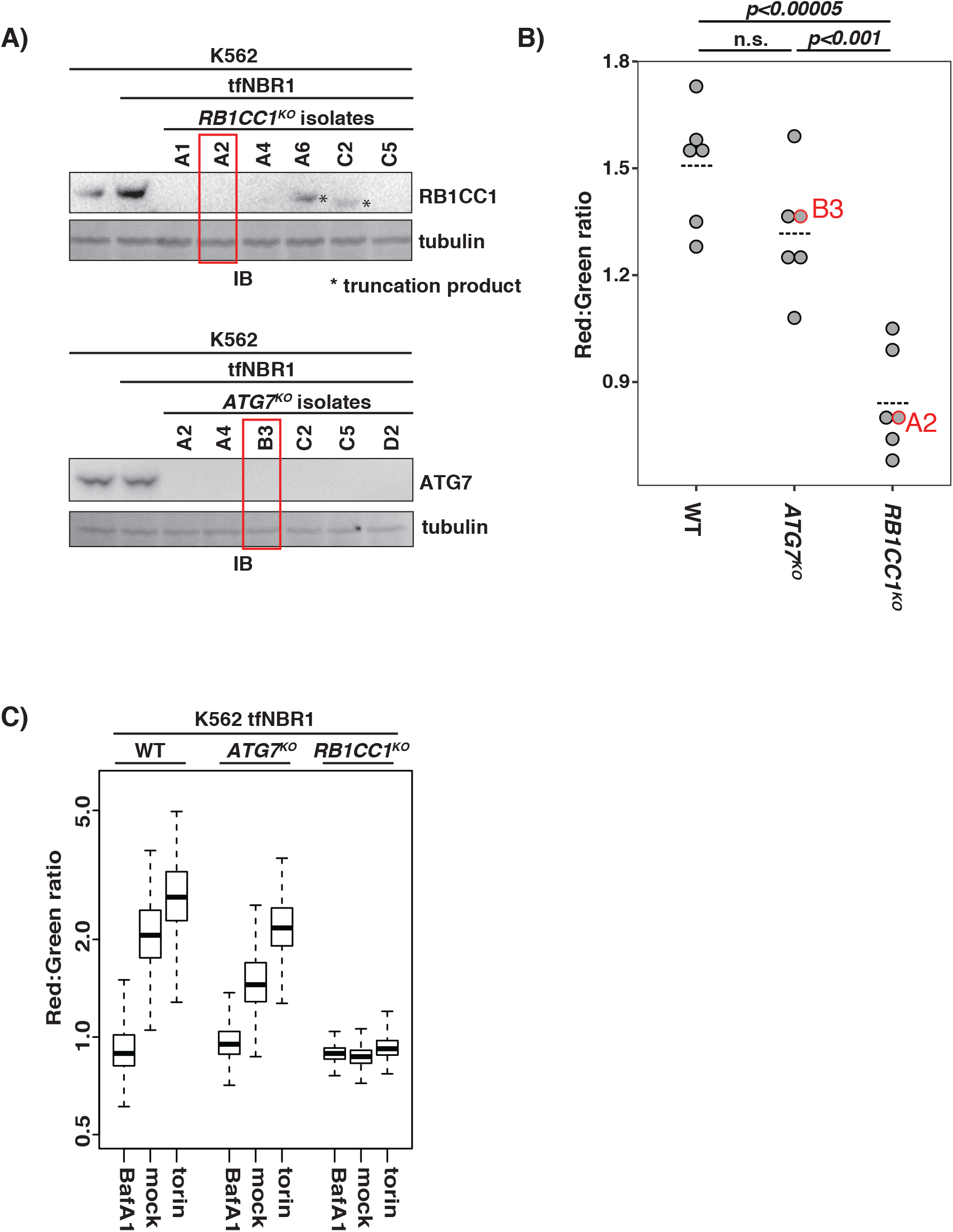
Figure 7, supplement 1. Comparison of *ATG7^KO^* and *RB1CC1^KO^* isolates. **(A)** K562 cells (far left lane) expressing tfNBR1 were transduced with sgRB1CC1 and sgATG7. Following clonal expansion of transductants, six candidates were selected for analysis by SDS-PAGE and immunoblotting (IB) of their cell extracts with indicated antibodies, as appropriate. Wildtype K562 cells and the parent tfNBR1 integrant cell line were also analyzed for comparison. Red boxes indicate isolates used in subsequent studies. **(B)** K562 tfNBR1 transductants described in part A and six WT clones were analyzed by flow cytometry (n>5000 cells). Shown are plots of median for individual isolates (dots) and the mean for the isolates combined (dashed lines). p-values (*ns*, not significant) shown were determined using a student’s t-test. Isolates used in subsequent studies are indicated in red. **(C)** K562 knockout isolates described in parts A and B and their wildtype parent were analyzed by flow cytometry under basal conditions and after 18h treatment with 250nM torin or 100nM BafilomycinA1 (BafA1). Plots show mean Red:Green ratios with inner quartiles (boxed regions) and top and bottom decade indicated (whiskers). n > 5000 cells each sample.

**Figure 2-source data 1. Read counts from all screens.**

Normalized read counts from each sequencing experiment. See ‘read_me’ tab for sample identity. Up, top 30% of cells by Red:Green ratio; Down, bottom 30% of cells by Red:Green ratio.

**Figure 2-source data 2. Beta scores from all screens.**

Individual beta scores from each experiment are listed by gene. Beta values from all replicates were averaged and are shown. Averaged values are color coded from red (suppressor) to white (neutral) to blue (enhancer). To view the underlying beta values for each replicate of a given reporter, expand the desired column by clicking the ‘+’ at the top the column.

Table S1. sgRNA oligos

List of 20-nucleotide oligos (plus overhangs) used to clone individual sgRNAs.

**Table S2. Other oligos**

List of oligos used in T7 endonuclease assays to confirm TMEM41B deletion and oligos used for endogenous labeling of TMEM41B with the 11^th^ beta strand of GFP.

**Table S3. List of primers for cloning tfReporters**

List of primers used to clone LC3B and receptors into the KpnI site of tfEmpty to generate each tfReporter.

**Table S4. Illumina primers for Brunello library**

List of primers used for Illumina sequencing of the Brunello library. Staggered oligos were pooled prior to PCR amplification. In contrast, a unique barcode primer was used for each sample. Green, Illumina P5 or P7 sequence; blue, sequencing primer annealing region; underline, stagger; red, region complementary to vector backbone (for PCR); purple, unique barcode (6-mer);

## Methods

### Antibodies

The following antibodies were used in this study. For immunoblotting (IB), all primary antibodies were used 1:1000 except where otherwise noted; secondary antibodies were used 1:3000 (HRP) or 1:10000 (fluorescent). For immunofluorescence (IF), antibody dilutions are noted following each antibody; secondary antibodies were used 1:500. Primary antibodies: mouse anti-SQSTM1 ([1:200 – IF] ab56416, Abcam), rabbit anti-TAX1BP1 (5105, CST), rabbit anti-NDP52 (9036, CST), rabbit anti-LC3B ([1:2000 – IB] NB100-2220, Novus), rabbit anti-LC3A/B ([1:100 – IF] 12741S, CST), rat anti-tubulin ([1:500 – IB] sc-53030, Santa Cruz), anti-p70-S6K (9202S,CST), anti-p70-S6K(pT389) (9205S, CST), rabbit anti-ATG13 (ABC344, EMDmillipore), rabbit anti-ATG13(pS318) (NBP2-19127, Novus), anti-WIPI2 ([1:150 – IF] ab105459, Abcam), rabbit anti-ATG7 (8558, CST), anti-RB1CC1 (12436S, CST), mouse anti-GFP (11814460001, Sigma), mouse anti-SERCA2 (ab2861, Abcam), rabbit anti-Calnexin (2679, CST); secondary antibodies (IB): goat anti-mouse IgG HRP (170-6516, Bio-rad), goat anti-rabbit IgG HRP (170-6515, Bio-rad), goat anti-rat IgG Alexa Fluor 488 (A11006, Invitrogen), goat anti-mouse IgG Cy5 (A10524, Invitrogen), goat anti-rabbit IgG Alexa Fluor 546 (A11010, Invitrogen); secondary antibodies (IF): goat anti-rabbit IgG Alexa Fluor 568 (A11036, Invitrogen), goat anti-mouse IgG Alexa Fluor Plus 488 (A32723, Invitrogen).

### Chemicals and reagents

The following chemicals and reagents were used in this study: torin1 (sc-396760, Santa Cruz), BafilomycinA1 (tlrl-baf1, Invivogen), poly-l-lysine (P4707-50ML, Sigma), Lipofectamine 3000 (L3000008, Life Technologies), nucleofector kit T (VACA-1002, Lonza), polybrene (H9268-5G, Sigma), normocin (ant-nr-1, Invivogen), puromycin (ant-pr-1, Invivogen), blasticidin (ant-bl-1, Invivogen), zeocin (ant-zn-1, Invivogen), normal goat serum (ab7481, Abcam), Phusion High-Fidelity DNA polymerase (M0530L, NEB), T5 exonuclease (M0363S, NEB), Taq DNA ligase (M0208L, NEB).

### Vectors

pHR-SFFV-GFP1-10 (Addgene plasmid # 80409) and pCDNA CMV mCherry-GFP-11 were gifts from Bo Huang. pMRXIP GFP-Stx17TM was a gift from Noboru Mizushima (Addgene plasmid # 45910). Human Brunello CRISPR knockout pooled library was a gift from David Root and John Doench (Addgene #73178). lentiCRISPRv2 puro was a gift from Brett Stringer (Addgene plasmid # 98290). lentiGuide-puro was a gift from Feng Zhang (Addgene plasmid # 52963). ptfLC3 was a gift from Tamotsu Yoshimori (Addgene plasmid # 21074). pX330-U6-Chimeric_BB-CBh-hSpCas9 was a gift from Feng Zhang (Addgene plasmid # 42230). AAVS1-CAG-hrGFP was a gift from Su-Chun Zhang (Addgene plasmid # 52344). psPAX2 was a gift from Didier Trono (Addgene plasmid # 12260). pCMV-VSV-G was a gift from Bob Weinberg (Addgene plasmid # 8454). pUMVC was a gift from Bob Weinberg (Addgene plasmid # 8449). pFUGW-EFSp-Cas9-P2A-Zeo (pAWp30) was a gift from Timothy Lu (Addgene plasmid # 73857).

### Isothermal assembly

PCR fragments were generated using 2X phusion master mix (M0531S, NEB) and insert-specific primers that appended a 30bp overlap with target DNA. Vector backbones were linearized by restriction enzyme and dephosphorylated by calf intestinal phosphatase (M0290S, NEB). Prior to assembly, all DNA fragments were gel purified (D4002, Zymo Research). 50ng of linearized vector DNA were combined with isomolar amounts of purified insert(s). 5ul of the resulting DNA mix was added to isothermal assembly master mix and incubated at 50 C for 20 minutes (Gibson et al., 2010). Assembled product was transformed into NEB Stable competent cells (C3040H, NEB) and plated on LB+agar plates (plus appropriate antibiotics) to isolate single isolates. Single isolates were grown in LB broth + antibiotics and plasmid DNA was purified using a Qiagen miniprep kit (27106, Qiagen). Sequences were verified by Sanger sequencing (Eton Bioscience Inc).

### sgRNA oligonucleotide (oligo) ligation protocol

sgRNA oligos were ordered from Eton Biosciences. Oligo sequences are listed in Table S1. To generate the necessary overhangs, all oligos were in the form: Forward: 5’ CACCGNNNNNNNNNNNNNNNNNNNN – 3’; Reverse: 5’-AAACNNNNNNNNNNNNNNNNNNNNC –3’. Oligos were diluted to 10uM in distilled water. 50pmol each of forward and reverse oligo were combined in a 25ul reaction and phosphorylated with T4 polynucleotide kinase (M0201S, NEB) in 1X T4 DNA ligase buffer (B0202S, NEB) for 30 minutes at 37 degrees Celsius. Phosphorylated oligos were boiled for 5 min at 95 degrees Celsius and slow cooled (0.1 degrees Celsiuslsec) to facilitate annealing. Annealed oligos were diluted 1:100 and 2μl of insert were ligated into 20ng digested vector (pLentiuGuide-puro, BsmBI site; pLentiCRIPSR v2, BbsI site; pX330-derivatives, BbsI site) using T4 DNA ligase (M0202S, NEB). Ligation was allowed to proceed for 15 minutes at room temperature. Ligated products were transformed into NEB Stable cells.

### T7 endonuclease (Surveyor) assay

Genomic DNA (gDNA) was extracted using QuickExtract buffer (QE0905T, Epicentre) according to the manufacturer’s instructions. gDNA was subsequently normalized to 200nglμl. 600ng gDNA was used per 100 μl PCR to amplify the targeted region of interest. Primers are listed in Table S2. The resulting PCR amplicon was purified using a Zymoclean Gel DNA Recovery Kit (D4002, Zymo Research) and normalized to 20ng/μl in 19ul 1X NEBuffer 2 (B7002S, NEB). Amplicon was boiled and cooled (-1 degrees Celsius/sec) to allow for hybridization. 1 μl T7 endonuclease I (M0302, NEB) was added and incubated for 15 min at 37 degrees Celsius. Reaction was quenched by adding 1.5μl 0.25M EDTA and analyzed in a 2% UltraPure Agarose gel (16500500, Invitrogen).

### Generation of AAVS1-targeting tfReporters

The hrGFP fragment was excised from AAVS1-CAG-hrGFP using SalI/EcoRV and replaced with RFP-GFP subcloned from ptfLC3 to generate pCS418 tfEmpty (puro). LC3B (NP_074058.2), SQSTM1 (NP_003891.1), NDP52 (NP_005890.2), NBR1 (NP_006015.4) or TAX1BP1 (NP_005822.1) were PCR amplified and subcloned into the KpnI site of pCS418. Primers are listed in Table S3. To generate blasticidin resistant versions of each cassette, the puromycin resistance gene was excised from pCS418 with an XhoI/SpeI digest and replaced with an analogous geneblock fragment (oVD5075, Integrated DNA technologies) encoding a blasticidin resistance gene (BSD).

### Tissue Culture

All cells were grown in a standard water-jacketed incubator with 5% CO2. K562 cells were grown in IMDM media (30-2005, ATCC) with 10% FBS (30-2020, ATCC) and 1x penicillin/strep. Cells were maintained below 1 million cells per milliliter. HEK293T cells were grown in DMEM media (30-2002, ATCC) with 10% FBS and 1x pen/strep. Normocin (1:500) was used as a common additive. All cells were passaged less than 25 times. For passaging, cells were trypsinized with Trypsin-EDTA (25300-054, Life Technologies). Puromycin (2μg/ml), blasticidin (5μg/ml) and zeocin (50μg/ml) were added when necessary for selection.

### Cell Line Authentifícation

Genomic DNA was isolated from HEK293T and K562 cells using the GenElute™ Mammalian Genomic DNA Miniprep Kits (Sigma-Aldrich). STR profiling and allele identification were performed by the Molecular Diagnostics Laboratory of Dana-Farber Cancer Institute. Briefly, isolated genomic DNA was analyzed with the GenePrint^®^ 10 short tandem repeat (STR) profiling kit (Promega) and Amelogenin for gender identification. GeneMapper v4 Fragment Analysis software (Applied Biosystems) and GenePrint10^®^ allele panel (Promega) custom bin files were used to identify the alleles at eight STR loci (TH01, TPOX, vWA, CSF1PO, D16S539, D7S820, D13S317 and D5S818). The ATCC STR Profile Database was used to verify that the identified alleles matched those of the expected cell type.

### Transient transfection and nucleofection

Prior to transfection, HEK293T cells were seeded in OptiMEM Reduced Serum media (51985-034, ThermoFisher). At 90% confluency, cells were transfected overnight using Lipofectamine 3000 reagent according to the manufacturer’s recommendations. The following morning media was exchanged and cells were passaged for another 24 hours prior to drug treatment. K562 cells were nucleofected using a Nucleofector™ 2b Device (Lonza) using nucleofector kit T (VACA-1002, Lonza) and protocol T-016. Transfectedlnucleofected DNA was prepared using a ZymoPURE™ Plasmid Midiprep Kit (D4200, Zymo Research).

Generation of gene knock-out cell lines using CRISPR-Cas9 gene editing: For the generation of stable knock-outs, HEK293T and K562 cells were transfected or nucleofected, respectively, as described above. Oligo sequences for sgRNAs were generated by CHOPCHOP or extracted from the Brunello library and cloned into the indicated vectors as outlined above under ‘sgRNA oligonucleotide (oligo) ligation protocol’ (Doench et al., 2016; Labun et al., 2016). Oligos are listed under Table S2. Cells were diluted by limiting dilution or cell sorting into 96 well plates and clonally expanded. Expanded cells were confirmed for knock-out by western blot or PCR/T7 endonuclease testing.

### Generation of GFP1-10 cell line

HEK293T cells were transduced with a lentiviral GFP1-10 expression cassette. Clonal cell lines were established by limiting dilution and validated for GFP1-10 insertion by transient transfection with mCherry-GFP-11. Four successful clones were saved and one was used for all further experiments. To tag TMEM41B, cells were transfected with pCS651 (coexpressing Cas9-T2A-BFP, sgTMEM41B) and oVD6217 (an oligo containing 5’homology arm-GFP11-linker-3’ homology arm). Successful integrants were identified and sorted by FACS.

### AAVS1 integration

Cell lines were co-transfected with pX330-U6-Chimeric_BB-CBh-hSpCas9-AAVS1-gRNA and AAVS1-tfReporter template vectors. 48 hours post-transfection cells were incubated with appropriate selection media and passaged for 14 days. Red+/Green+ cells were sorted and propagated.

### Lentiviral generation

Lentivirus was generated in HEK293T cells using Lipofectamine 3000. Cells were grown overnight in Opti-MEM media (5% FBS, no antibiotics) (31985062, ThermoFisher) to 90% confluency. Cells were than transfected with pVSV-G, pSPAX2 and packaging constructs at a 1:3:4 ratio. Transfection proceeded for 6-8 hour before media was refreshed. Virus was collected and pooled at 24 and 48 hours post-transfection. Virus was pelleted at 1000g 2X 10 min, aliquoted and frozen in single-use aliquots. For retrovirus production, all methods were the same except pUMVC was exchanged for pSPAX2.

### Viral transduction

Cells were incubated in appropriate media containing 8μglml polybrene and lacking penicillin/streptomycin. Cells were transduced overnight. Media was exchanged for media lacking polybrene for 24 hours prior to antibiotic selection.

### Protease protection assay

Our protocol for mammalian protease protection assay was based on Zhao et al. (Zhao et al., 2013). Cells were seeded so they would be 70% confluent at 5pm. Media was then exchanged into media containing 100nM Bafilomycin A1. Cells were incubated for 15 hours. After incubation, cells were trypsinized and pelleted. Cells were washed 1X with cold PBS and resuspended in pre-chilled lysis buffer (20mM Hepes KOH pH7.4, 0.22M mannitol, 0.07M sucrose). Cells were lysed by extrusion through a 30 gauge needle 10-15 times. Samples were pelleted 2X at 500g for 10 min at 4 degrees Celsius to pellet debris. When indicated, samples were incubated with 1X trypsin (T1426-100MG, Sigma; 100X stock: 1mglml) andlor 0. 5% Triton X-100 for 35 minutes at 30C. Reactions were quenched in 1X hot Laemmli sample buffer and boiled at 65 degrees for 10 minutes.

### Immunoprecipitation

Cells were collected and resuspended in IP buffer (50mM HEPES pH 7.4, 150mM NaCl, 2mM EDTA, 1% Triton X-100). Cells were incubated on ice for 30 minutes and pelleted twice at 5000g for 5 minutes at 4C. Supernatant was applied to pre-washed GFP-Trap or RFP-Trap magnetic agarose (Chromotek) and incubated for 1 hour at 4C. Beads were washed 4x 5 min with two tube changes. Protein was eluded by boiling at 70C in 1X SDS buffer.

### Gel electrophoresis and western blotting

Cells were trypsinized at 75% confluency and quenched in an equal amount of media. Cells were lysed for 15 minutes on ice in lysis buffer (50mM HEPES pH 7.4, 150mM NaCl, 2mM EDTA, 1% Triton X-100, 2X cOmplete protease inhibitor tablet [Roche]). For phosphorylation analysis, lysis buffer was supplemented with phosphatase inhibitors (10X: 100mM NaF, 10mM Na3VO4, 100mM NaPPi). Lysates were cleared 2X at 1000g for 5 minutes. Post-spin supernatants were used as input. Protein levels in supernatants were normalized using a BCA protein assay (Pierce #23227). Normalized samples were boiled in 1X (final concentration) Laemmli Loading Buffer (3X stock: 189 mM Tris pH6.8, 30% glycerol, 6% SDS, 10% beta-mercaptoethanol, bromophenol blue). Gel electrophoresis was performed at 195V for 70 minutes in Novex 4-20% Tris-Glycine gels. Total protein analysis was performed using SYPRO Ruby (Thermo Fisher) according the manufacturer’s recommendations (short protocol). For western blotting, samples were transferred for 60 minutes to 0.2μm PVDF membranes (Millipore Sigma #ISEQ00010) using a Semi-dry transfer cell (Bio-Rad). Membranes were blocked for 20 minutes in TBS-T with 5% milk. Primary antibodies were incubated overnight at 4 degrees Celsius. Blots were then rinse 3X 5 minutes. Secondary antibodies were incubated for 1 hour at room temperature. Blots were rinsed 4X 10 minutes in TBS-T and imaged using fluorescence (Typhoon Trio Imager) or chemiluminescence (SuperSignal™ West Femto Maximum Sensitivity Substrate, ThermoFisher Scientific, #34095). If necessary, stripping of membranes was performed using Restore Western Blot Stripping Buffer (ThermoFisher Scientific #21059) for 10 minutes.

### Immunofluorescence/Immunocytochemistry

Coverslips (Fisher Scientific, 12-548A) were placed in 6-well tissue culture plates (VWR, 62406-161) and coated with poly-L-lysine (Sigma, P4707) per the manufacturer’s recommendations. Cells were seeded onto coverslips overnight so that they would be 15% confluent at the time of fixation. When reported, cells were treated with 250nM torin for 1 to 3 hours prior to fixation. Fresh 16% PFA (Electron Microscopy Sciences, #15710) was diluted to 4% in 1X Dulbecco’s phosphate buffered saline with calcium chloride and magnesium chloride (10X: Life Technologies, #14080-055). Coverslips were removed with forceps and placed into 4% PFA for 15 minutes. PFA was aspirated and washed twice with PBS (Sigma, D8537). For LC3 immunofluroescence, cells were transferred to wells containing prechilled (-20C) methanol for 5 minutes. Slides were returned to PBS and washed 2X for 5 minutes. Slides were blocked at RT for 1 hour in blocking buffer (0.3% Triton X-100, 5% Normal Goat Serum (NGS) in PBS) and washed once in PBS. Primary antibody was diluted in 5% NGS at the dilutions described elsewhere in these methods. 75ul of antibody mixture was spotted on parafilm in a humidified chamber and inverted coverslips were incubated with antibody overnight at 4C. After incubation, coverslips were washed 3X 10 min in PBS. Secondary antibodies were diluted 1:500 in 5% NGS and inverted coverslips were incubated with antibody mixture for 45 minutes. Cells were stained with a 1:10,000 dilution of Hoechst 33342 (ThermoFisher Scientific, H3570) for 5 minutes. Coverslips were washed 4X 10 minutes in PBS and mounted on coverslips (Corning, 294875X25) using Prolong Diamond (ThermoFisher Scientific, P36965).

### Confocal microscopy

Fluorescent images were obtained using a confocal microscope with Airyscan detectors (LSM880 with Airyscan, Zeiss), a 63x PlanAPO oil-immersion objective lens (Zeiss), and processed with Zeiss Blue software (Zeiss).

### Image analysis

Fluorescence microscopy images were processed using a newly developed Python analysis pipeline built around the pyto_segmenter analysis package (Weir et al., 2017). First, regions of images containing cells were identified. To do so we first fit a Gaussian distribution to the fluorescence intensity distribution for a smoothed green channel (488 nm excitation) z-stack from an empty field. Using this Gaussian fit we predicted the probability that each pixel in the smoothed green channel z-stack for each field corresponded to background (noncell) or foreground (cell). We assigned each pixel with a p(background) < 10^−5^ to the cells, thus creating a “cell mask”. After removing small specks (<100,000 pixels volume) to eliminate debris, we removed out-of-focus planes from the cell mask using a Support Vector Machine (SVM) classifier as described previously (Hsu and Chen, 2008). Next, nuclei were segmented from the blue (DAPI) channel by slice-by-slice relative thresholding followed by watershed segmentation using the pyto_segmenter package. Cells were segmented using watershed segmentation from nuclei seeds. Cell edges were eroded to eliminate blurred edge excess included during the p-value transformation. Cells contacting the edge of the field were removed from analysis. Next, punctae were segmented in the green and red (561 nm) channels using the pyto_segmenter package with empirically determined Canny edge detection thresholds. The number of total punctae and punctae overlapping with objects in the other fluorescence channel were counted and tabulated data were saved in .csv format. Plotting was performed using R and the ggplot2 package. See the image analysis package for details. Scripts for image analysis, scripts for data plotting and tabulated values from microscopy data can be found at <u>https://github.com/deniclab/csth-imaging/tree/pubversion</u>.

### Sample size estimate and experimental replication details

For microscopy experiments, 40 images were collected and the number of cells in each sample was counted using segmentation scripts. Cell counts are indicated above each quantitation. Samples were masked prior to data collection and analyzed using automated scripts to eliminate bias during quantification. Replicates represent biological replicates in which strains were subjected to identical preparations on different days.

For sequencing experiments, the number of replicates (2-4) are indicted in Figure 2-source data 1. Each replicate was a biological replicate where strains were transfected and taken through the entire experiment on separate days.

For flow cytometry experiments, N is indicated for each experiment in the figure legend. >5000 cells were used for each experiment.

## Library propagation

Brunello library (2 vector system) was purchased from addgene (item #73178). 50ng of library was electroporated into 25 μl Endura electrocompetent cells (Lucigen, 60242-2). Cells from 8 electroporations were pooled and rescued in 8 ml of rescue media for 1 hour at 37 degrees Celsius. 8 ml of SOC (2% tryptone, 0.5% yeast extract, 10 mM NaCl, 2.5 mM KCl, 10 mM MgCl2, 10 mM MgSO4, and 20 mM glucose) was added to cells and 200μl of the final solution was spread onto 10 cm LB plates containing 50 μglml carbenicillin (80 plates total). Through a dilution series, 500 million colonies were estimated, representing 7000x coverage of the library. Cells were manually scraped off plates and a GenElute Megaprep kit (Sigma, NA0600-1KT) was used to purify plasmid DNA.

## Library lentiviral generation

Lentivirus was generated by lipofection (Lipofectamine 3000) of HEK293T cells with 5 μg psPAX2 (Addgene Plasmid #12260), 1.33 μg pCMV-VSV-G (Addgene plasmid Plasmid #8454), and 4 μg library vector per 10 cm plate. Transfection was performed according to manufacturer’s specifications. Briefly, low-passage HEK293T cells were grown in Opti-MEM + 5% FBS medium to 95% confluency by time of transfection. Cells were transfected for 6 hours, then media was replaced with fresh Opti-MEM + 5% FBS. 24 hours post-transfection, supernatant was collected and replaced. 48 hours post-transfection, supernatant was again collected, pooled with the 24 hour supernatant and clarified 2X 1000g for 10 min. Viral RNA was purified using a Machery Nagel viral RNA purification kit. Viral RNA was quantified using the Lenti-X™ qRT-PCR Titration Kit (Takara). A value of 849 copieslIFU, derived from a control virus expressing BFP, was used to calculate viral titer.

## Transduction and cell growth

For CRISPR screening experiments, K562 cells were passaged to maintain cell density between 500,000 and 2 million cellslml. Cells were propagated in IMDM + 10% FBS + penlstrep + appropriate antibiotics (Blasticidin 5 μglml, zeocin 50 μglml) until 200 million cells were obtained (approximately 8-10 days). All this stage, all remaining experiments were performed in 175 cm^2^ flasks. For infection, 200 million cells were pelleted and resuspended in IMDM + 10% FBS + 8 μglml polybrene. Date of infection was day 0. An MOI of 0.4 was used to minimize multiple infection events per cell. Cells were infected overnight, pelleted and exchanged into fresh media. After 24 hours, cells were split and 2 μglml puromycin was added. Cells were continually passaged in puromycin. At day 10, cells were removed from puromycin and at day 12 cells were sorted for red:green fluorescence. 100M unsorted cells were pelleted and processed as input. The top and bottom 30% of cells (based on Red:Green ratio) were taken. 100 million cells were sorted for each experimental condition. Cell sorting was performed using a FACSAria or BioRad S3 sorter. Cells were pelleted and stored at -80 degrees Celsius until processing.

## CRISPR screen processing

Genomic DNA was purified from collected cells using the NucleoSpin Blood XL kit (740950.1, Machery Nagel) according to the manufacturer’s instructions. Illumina sequencing libraries were created by PCR amplifying the genomically-integrated sgRNA sequences. All genomic DNA was used for each PCR. A pool of eight staggered-length forward primers was used in each PCR reaction to avoid monotemplating during Illumina sequencing. Reverse primers contained unique barcodes designed to allow for sequencing and differentiation of multiple libraries on a single chip during Illumina sequencing. Each 50 μL PCR reaction contained 0.4 μM of each forward and reverse primer mix (Integrated DNA Technologies), 1x Phusion HF Reaction Buffer (NEB), 0.2 mM dNTPs (NEB), 40 U/mL Phusion HF DNA Polymerase (NEB), 5 μg of genomic DNA, and 3% v/v DMSO. The following PCR cycling conditions were used: 1x 98°C for 30 s; 25x (98°C for 30 s, 56°C for 30 s, 72°C for 30 s); 1x 72 °C for 10 min. The resulting products were pooled to obtain the sgRNA libraries. The pooled PCR products were size selected by adding 0.95x magnetic bead slurry as outlined by DeAngelis et al.(DeAngelis et al., 1995). The High Sensitivity D1000 ScreenTape system (Agilent Technologies) was used to confirm the absence of primer dimers after purification. Sample libraries were quantified by qPCR using the NEBNext Library Quant Kit for Illumina (NEB). Four to five libraries were pooled to a total concentration of 10 nM for simultaneous Illumina sequencing on a single chip. The libraries were sequenced using either the HiSeq 2000 or 2500 system (Illumina). The HiSeq 2000 system was run on High Output Run Mode with the TruSeq SBS V3 kit (Illumina). The HiSeq 2500 system was run on either the Rapid Run Mode with the HiSeq Rapid SBS V2 kit (Illumina) or the High Output Run Mode with the HiSeq SBS V4 kit (Illumina). Sequencing was performed per recommendations of the manufacturer with custom sequencing and indexing primers (Integrated DNA Technologies). For primer sequences, see Table S4.

## NGS data analysis

The 5’ end of NGS reads were trimmed to 5’-CACCG-3’ using Cutadapt. The count function of MAGeCK (version 0.5.3) was used to extract read counts for each sgRNA. The mle function was used to compare read counts from cells displaying increased and decreased Red:Green ratios.

The output included both beta scores and false discovery rates. Beta scores for each sgRNA for each tfReporter were averaged across 2-4 experiments. Across all experiments, average read counts were 200-400 per sgRNA. To generate heatmaps for each reporter (e.g. Fig 2), the beta scores for each gene were normalized by the beta score for ATG9A.

## Flow cytometry

All samples were pelleted, washed 1X in cold PBS, and filtered through strainer cap tubes (VWR, 21008-948) prior to analysis. Flow cytometry data were collected on an LSRII flow cytometer. Data were analyzed in FlowJo (FlowJo, LLC) and R. The biomodality of populations was determined in an unbiased manner using the BifurGate tool in FlowJo. At least 5000 cells were collected for all samples.

